# Defective neuritogenesis in *Abcd1/2* deficient rat neurons due to intrinsic and astrocyte-dependent mechanisms

**DOI:** 10.1101/2022.09.30.510337

**Authors:** Arantxa Golbano, Luis Pardo, Carmen M. Menacho, Marina Rierola, Enrique Claro, Levi B. Wood, Roser Masgrau, Elena Galea

## Abstract

X-linked adrenoleukodystrophy (X-ALD) is a rare neurometabolic and demyelinating disorder caused by loss of function mutations of the ABCD1 transporter that imports very-long-chain fatty acids (VLCFA) into the peroxisome for beta-oxidation. Impaired ABCD1 function results in VLCFA accumulation, which ultimately causes lethal forms of X-ALD in children (CCALD) and adults (CAMN). Because X-ALD is a genetic disorder, we looked for signs of altered neurodevelopmental pathways in the transcriptomes of brain cortical tissues free of pathology from patients that died of CALD or CAMN. Several categories related to brain development, axonal growth, synaptic signaling and synaptic compartments were significantly dysregulated in both CALD and CAMN, suggesting that congenital circuit abnormalities might be structural in brains of mutated ABCD1 carriers. We partially dissected the cellular origin of dysregulated pathways using rat neuronal and astrocytic cultures in which X-ALD was modeled by silencing of Abcd1 and Abcd2 by RNA interference. Abcd2 was silenced lest it compensated for Abcd1 loss. Abcd1/2 deficient neurons presented higher rates of death, reduced sizes and defective formation of spines, dendrites and axons. The aberrant neuron development was caused by cell-autonomous and astrocyte-dependent mechanisms, and involved Wnt signaling, as suggested by the rescue of the expression of a synaptic gene upon pharmacological activation of the Wnt pathway. As recently proposed for neurogenetic disorders such as Huntington’s disease, our data suggest that X-ALD has a neurodevelopmental component that may cause psychiatric alterations and prime neural circuits for neurodegeneration. If this is the case, therapies aimed at restoring neural-circuit function in neurodevelopmental disorders may be reprofiled for X-ALD therapeutics.

## Introduction

X-linked adrenoleukodystrophy (X-ALD) is a rare neurodegenerative disorder affecting 1 in 17000 births, mostly men^1^. It is caused by loss-of-function mutations of a gene encoding the ATP Binding Cassette Subfamily D type 1 (ABCD1) peroxisomal transporter, which imports very-long-chain fatty acids (VLCFA) (i.e., saturated fatty acids 22-carbons-long or more) into the peroxisome for degradation by β-oxidation^2^. Mutated *ABCD1* genes generate non-functional or unstable proteins that are readily eliminated, causing the aberrant accumulation of VLCFA in plasma and tissues, such as the adrenal glands, the CNS and the testis^3^.

VLCFA in high concentrations are toxic to neurons, oligodendrocytes, microglia and astrocytes^4–8^. How this cellular damage leads to X-ALD is an outstanding question, for the disease, of which over 900 variants have been documented^9^, manifests itself in a phenotypic spectrum that ranges from adrenal insufficiency to fatal cerebral demyelinating disease. The absence of genotype-phenotype correlations represents a major complication in the clinical management of X-ALD, partially alleviated by a broad classification of patients according to their age at disease onset. In childhood X-ALD, called CCALD (double C for ‘childhood’ and ‘cerebral’), symptoms appear on average at 7 years of age and consist of intellectual, visual and gait disturbances^10^. CCALD is usually lethal^10^. When analyzed *postmortem*, the brains from these patients present extensive demyelination, microgliosis, astrogliosis, and perivascular lymphocytic infiltration^11,12^. In adults, mean disease onset is 30 years of age in men^10^, gait disturbance being the initial symptom^10^. Virtually all these patients and over 80% of the women develop a chronic form of X-ALD, referred to as adrenomyeloneuropathy (AMN), featuring progressive Wallerian degeneration of corticospinal tracts, peripheral sensory-motor neuropathy and adrenal insufficiency, with no overt brain demyelination^13,14^. Around 35-50% of male patients with AMN develop the lethal cerebral form of X-ALD, known as CAMN^10,15^.

Who, among individuals harboring *ABCD* mutations, will develop X-ALD, when, and with what severity, is unpredictable. Although disease modifiers^16–18^ and life hazards^19^ may determine X-ALD onset and progression, the molecular and cellular underpinnings of the phenotypic variability of X-ALD remain largely unknown. Nor is the cascade of pathological events driving CCALD, AMN and CAMN completely elucidated. Thus, whether axon injury in cerebral and spinal-cord manifestations of X-ALD is independent of, or secondary to, axon demyelination^20^; whether axonal damage is due to cell-autonomous VLCFA-elicited mitochondrial damage and oxidative stress, long axons being particularly vulnerable, as suggested for AMN based on mouse models^21^; whether microglia contribute to neuronal damage by loss-^7,22^ or gain-of-toxic function^23^, and, finally, what the contribution of dysfunctional astrocytes is^24,25^, are open questions.

Here, we asked whether X-ALD may alter neurodevelopment thereby rendering neural circuits more susceptible to neurodegeneration over time. The question is motivated by recent studies in Huntington disease (HD), a dominantly inherited disorder caused by a CAG expansion mutation in the huntingtin (HTT) gene. Human fetuses and mouse embryos with the HTT mutation show abnormalities in the developing cortex consistent with defective migration of progenitor cells^26^. Likewise, neurons^27^ and neural stem cells derived from induced pluripotent stem cells obtained from patients with HD^28^ show changes in gene expression pinpointing altered neurodevelopment. A striking discovery was the rescue of cognitive impairment and synaptic pathology upon pharmacological restoration of neurodevelopmental pathways in adult HD mice^27,28^. In causally linking neurodevelopmental alterations with pathology in adulthood, this evidence supports a rather counterintuitive idea: the therapeutic manipulation of altered neurodevelopmental pathways may still show benefits if performed in adult life. HD is indeed not the first disease for which an association between neurodegeneration and aberrant neurodevelopment has been proposed. Others are autism spectrum disorder, Fragile X syndrome, Alzheimer’s disease (particularly ApoE4 carriers) and Down syndrome^29^.

Arguably, X-ALD has not been considered a neurodevelopmental disorder because no gross anatomical or cytoarchitectural abnormalities characteristic of other peroxisomal disorders have been detected in brains of patients with X-ALD^30^, nor are there cognitive deficits in asymptomatic children carrying mutated ABCD1^31^. However, mild neuropsychiatric alterations or below average performances in cognitive tests have been reported in AMN patients with no cerebral pathology according to MRI^34–36^. Since X-ALD is a genetic disorder, we posit that the mild mental problems are due to structural deficits in neural circuits owing to ABCD1 malfunction and lipid dyshomeostasis during brain development.

Herein we searched for signs of altered neurodevelopment in X-ALD with a top-down approach consisting in using patient transcriptomic data to inform studies in cellular models of X-ALD. First, we looked for dysregulated pathways related to neuronal development in publicly available microarrays from non-diseased white matter cortical tissue of CCALD and CAMN patients^38^. We reasoned that the analysis of non-affected areas would inform about latent pathology. Second, since cellular compartmentalization is lost in transcriptomic analyses of whole tissue, we partially dissected out the cellular origin of altered neurodevelopmental pathways in cultures of neonatal rat brain neurons and astrocytes, in which we induced X-ALD-like conditions by silencing *Abcd1* and *Abcd2* transporters, as previously done in astrocytes^24,39^. Both transporters were silenced to model the most severe consequences of aberrant VLCFA accumulation, because *Abcd2* can partially compensate for the loss of *Abcd1*^40^. Since, obviously, the compensation by *ABCD2* is not completely effective in carriers of mutated *ABDC1* who develop X-ALD, we aim to model this scenario in rats by deleting *Abcd2*, too.

We report that the silencing of *Abcd1 and Abcd2* in neurons leads to aberrant neuronal growth, neuritogenesis, spinogenesis and axonogenesis due to a predominant, cell-autonomous mechanism, exacerbated by a secondary mechanism mediated by defective astrocyte conditioned media (ACM). Deficits in Wnt signaling might underlie the alterations in neurons and astrocyte-neuron interactions according to pharmacological and molecular analyses. The data support that a hitherto overlooked neurodevelopmental component may exist in X-ALD. This discovery might inform therapies for psychiatric symptoms, particularly in adult X-ALD, and perhaps help arrest the progression of AMN into CAMN by restoring neural-circuit homeostasis.

## Material and Methods

### Microarray data

Microarray raw data from X-ALD patients was obtained from the dataset with accession number E-MEXP-3288). The data had been obtained from 24 subjects (three CCALD, three control children, eight CAMN and ten adult controls) as described in^38^. The program available at arrayanalysis.org was used to generate fold change (FC), logFC, p-values and adjusted p-values of the comparisons between CCALD and CAMN groups with their respective controls (Suppl. data 1.) For CCALD vs controls, there were 3683 differentially expressed genes (DEG) (p < 0.05), of which 34% were upregulated and 65.4% downregulated. For CAMN vs controls, there were 5394 DEG (p < 0.05), 37.4% upregulated and 62.5% downregulated.

### Neuronal and astrocytic cultures

Neuronal and astrocytic cultures were obtained from neonatal Sprague-Dawley rat pups (0-1 days old) as previously described^41,42^. The manipulation of rats was performed according to CEEAH/UAB, Catalonian, Spanish and EU regulations. For each brain, the hippocampi were processed to generate neurons, and the cortex to produce astrocytes and ACM. Three reasons justify the use of two brain regions. First, reduction of animal numbers, since different parts of the same brain were used. Second, because hippocampal neurons are less mature than cortical ones at birth, hippocampus is preferable over cortex to study neuritogenesis postnatally. By contrast – and third reason – cortices are preferable over hippocampi to generate astrocytes because they are a much larger region, thereby ensuring sufficient ACM production to treat neuronal cultures.

The astrocytes were cultured in DMEM (Thermo Fisher) supplemented with 10% fetal bovine serum (FBS), 20 U mL^-1^ penicillin, and 20 μg mL^-1^ streptomycin, at 37°C in a humidified atmosphere containing 5% CO2. Upon reaching confluency (10 days), the cultures were agitated in a mechanical shaker to remove the microglia growing on top of the astrocytes. The astrocyte cultures were then trypsinized and the cells seeded on fresh flasks to generate secondary astrocyte cultures, which were the ones used herein. Cell composition was assessed with cell specific markers for astrocytes (GFAP), microglia (IBA1), oligodendrocytes (GALC) and neurons (NEUN). Astrocyte cultures were composed of 94.3% astrocytes and 5.4% microglial cells.

Hippocampal neurons were cultured in standard Neurobasal A (NBA) medium containing 1X serum-free B27 supplement, 1X GlutaMAX^TM^ and 20 U mL^-1^ penicillin and 20 μg mL^-1^ streptomycin at 37°C in a humidified atmosphere containing 5% CO2. Neuronal cultures were composed of 74.4% neurons, 17.4% astrocytes, 3.4% microglia and 4.8% oligodendrocytes (data not shown). Days in culture are indicated in

### Abcd1 and Abcd2 silencing

In astrocytes, the expression of *Abcd* transporters was silenced using synthetic siRNA for *Abcd1* (Ambion, s170867) and *Abcd2* (Ambion, s136486) (Supp. data 2). Secondary astrocyte cultures were transfected with the siRNAs using Lipofectamine® RNAiMAX (Invitrogen) in DMEM with no FBS and antibiotics to a final concentration of 20 nM. The Silencer® Select Negative Control (Ambion, Catalog # 489043) was used as a control of silencing specificity. The transfections were performed following the manufacturer instructions. The media were replaced after 4 h. Forty-eight h thereafter, the extent of silencing was assessed by mRNA quantification with qPCR.

Because the transfection of neurons with nucleic acid materials is not very efficient, the expression of *Abcd1* and *Abcd2* was suppressed in these cells using short-hairpin (sh)RNA technology instead of siRNA. The sense and antisense sequences of the shRNAs for *Abcd1* and *Abcd2* were the same as the ones used for siRNA in astrocytes, and were connected with a loop replacing two timidines at the end of the sense sequence. A scrambled shRNA was used as a negative control. The shRNAs were delivered to neurons by adenovirus type 5 viral vectors (Ad5) generated by the Viral Vector Production Unit of the Universitat Autònoma de Barcelona (UPV-UAB). Before the infections, 50% of the media was removed and kept aside. Viral-vector particles were left overnight. The next day, the media was replaced with 50% of fresh media and 50% of conditioned media.

Ad5-shRNAs for *Abcd1* and *Abcd2* were tested alone or in combination at different multiplicity of infections (MOI, 10-2000). A MOI of 50 showed the highest silencing capacity. Thus, neurons were co-infected with the shRNAs for *Abcd1* and *Abcd2* at a MOI 50 each or with shScrambled at a MOI 100. Viral-vector infections were performed at 0 or 7 days in vitro (DIV) for short- and long-term studies, as indicated for each experiment.

### Treatments

#### Hexacosanoic acid (C26:0)

C26:0 (Sigma-Aldrich) was used to exacerbate the X-ALD phenotypes because it is the most abundant VLCFA in the plasma of patients^43^. C26:0 was dissolved in the synthetic carrier α-cyclodextrin (α-cyclo, Sigma Aldrich). A stock of C26:0 was prepared in chloroform and stored at −80°C. Before use, the chloroform was evaporated with gas nitrogen, and the resultant powder was dissolved in 2.5 μM α-cyclo and was sonicated for 20 s at a 30% power. Neuronal and astrocyte cultures were treated for 96 h with 50 μM C26:0.

#### ACM production

Control or *Abcd1/2* null astrocytes were subjected to a 4-h pulse of 100 μM C26:0 or vehicle in DMEM without glucose containing 1% FBS, 20 U/mL-penicillin and 20 μg/mL streptomycin. To make ACM compatible with neurons, DMEM was then replaced by the neuronal media NBA containing 1X serum-free B27 supplement minus antioxidants (minus AO, ThermoFisher), 1x GlutaMax (ThermoFisher), 20 U/mL penicillin and 20 μg/mL streptomycin, and left 48 h. Since oxidative stress contributes to X-ALD^44^, NBA minus AO and not regular NBA was used to produce ACM lest the damage inflicted by *Abcd* depletion and C26:0 in neurons be mitigated by antioxidants present in ACM. ACM were stored at −80°C.

#### GSK-3 inhibition

ShScrambled*/*α-cyclo and sh*Abcd/*C26:0 neurons were treated every 24 h for 96 h with 5 μM of the selective GSK inhibitor SB216763 (Sigma Aldrich; S3442) dissolved in dimethyl sulfoxide (DMSO). DMSO was used as a control. The cells were processed to obtain mRNA for the assessment of *Syn* and *Gapdh* expression by qPCR.

### Immunofluorescence

The cells were fixed with 4% paraformaldehyde (Electron Microscopy Science; 15714) for 15 min at room temperature (RT). The cells were permeabilized using 0.1% Triton X-100 (Alfa Aesar; A16046) diluted in PBS for 15 minutes, followed by a 30-min blockade with 5% normal goat serum (NGS) (GibcoTM; 16210-064) in PBS. To immunostain neurofilament SMI 312, as it is a phosphorylated protein, 0.02% saponin was used to permeabilize cells, and 1X TBS was the general buffer instead of PBS. The cells were incubated overnight with anti-NEUN (1/1000; Merck Millipore MAB377), anti-GFAP (1/1000; Dako Z0334), anti-GALC (1/500; Merck Millipore AB142), anti-IBA1 (1/1000; Wako 019-19741), anti-Vimentin (1/500; Abcam ab8978), anti-neurofilament marker SMI 312 (1/500; BioLegend 837904), all diluted in blocking solution. The next day, the cells were incubated with the corresponding secondary antibodies, either goat anti-mouse IgG 594 (1/1000; Thermo Fisher Scientific A-11032) or goat anti-rabbit IgG 488 (1/1000; Thermo Fisher Scientific A-11034) for 1 h at RT and with DAPI (1/20,000; Sigma) for 5 min at RT. The coverslips were mounted on slides using Fluoromont-G (Southern Biotech 0100-00.) F-actin was stained with the ActinGreen^TM^ 488 (ReadyProbes^TM^.)

### Cell death assays

Neuronal and astrocytic death was measured with the terminal deoxynucleotidyl transferase (TdT) dUTP Nick-End Labeling (TUNEL) Apo-Green kit from Biotool on fixed cells. Co-localization with cell-specific markers was determined by immunofluorescence as described above.

### Image analysis

The images to assess the cellular composition of cultures and cell death were acquired with a fluorescence microscope Nikon Eclipse TE2000-E connected to a CCD EG-ORCA camera (Hamamatsu Photonics) using Metamorph software (Universal Imaging). The plug-in ‘cell counter’ of Fiji/ImageJ software was used to count TUNEL positive cells. Images for the assessment of neuritogenesis were obtained with a confocal microscope Zeiss LSM700. Fluorescence was transformed into tridimensional (3D) masks with the ‘filament tracer’ plug-in of the IMARIS software (Bitplane, South Windsor, CT, USA) to quantify spine formation, dendrite arborization and thickness and axonal growth. To generate masks, neuronal somata were labeled with 20-μm diameter spheres. Seed points were placed along neurites starting 40 μm away from the somata center of mass. The minimum dendrite thickness was set at 0.35 μm, the minimum width detectable. The minimum and maximum spine diameters were set at 0.5 and 1.5 μm, respectively, for they were the smallest and largest widths detected. Once established, the same parameters were used to create masks from all fluorescence signals. Data per condition were generated from at least 15 images from each independent experiment.

### Western blots

Protein extraction and western blots were performed as described^38^. The primary rabbit monoclonal anti-ABCD1/ALD antibody (ab197013, Sigma Aldrich, 1/1000) was incubated overnight at 4°C. The primary antibody mouse monoclonal anti-β-ACTIN (a5316; Sigma Aldrich; 1/20000), used as a loading control, was incubated for 45 minutes at RT.

### Quantitative PCR

Total RNAs were isolated using the Trizol^TM^ reagent and were reverse transcribed as described before^45^. The samples were diluted and the cDNAs quantified using SYBR green (Life Technologies) in the case of *Syn* and *Gapdh*, or with Taqman probes for the rest of the genes, using in all cases a 7500 Fast Real-Time PCR System (Applied Biosystems). *Gapdh* was the house-keeping gene used to normalized samples. Data analysis was performed with the comparative Cq method using LingReg software^45^. Information about probes is in Suppl. Data 3.

### Measurement of cholesterol biosynthesis, accumulation and secretion

Cholesterol synthesis was measured in astrocytes by determining the conversion of the cholesterol precursor ^14^C-acetate into ^14^C-cholesterol. *siScrambled* or *siAbdc1/2* Astrocytes were incubated overnight with DMEM without glucose supplemented with 1% FBS containing 1 μCi/mL ^14^C-Acetate ([1-^14^C] acetate, 58 mCi/mmol, PerkinElmer) and or α-cyclo or 50 μM C26:0. The astrocytes were washed with PBS and processed for the extraction and separation of lipids. Chloroform/methanol (1:2, v/v) was added to solubilize lipids. 0.6 mL were transferred to a polypropylene tube and mixed with 0.5 mL of chloroform and 0.5 mL of H2O. The tubes were vigorously agitated and centrifuged for 5 minutes at 1000 rpm. Two phases appeared after the centrifugation, an aqueous phase (superior) and an organic phase (inferior). The organic phase containing chloroform was collected and transferred to new tubes. The chloroform was evaporated using a speed-vac for 1 h. The lipidic bottoms were resuspended in 10 μL of chloroform/methanol 1:1, then 10 μg unlabeled cholesterol was added as carrier, and samples were spotted onto the concentration zone of high performance thin-layer chromatography (HPTLC) plates (Merck). The chromatography plates were developed with hexane/ethyl ether/acetic acid (70:30:1, v/v/v), air-dried and sprayed with primuline (5 mg in 100 mL acetone / H20 (80:20, v/v). The bands were visualized with UV, marked with a pencil, scraped from the glass base, and transferred to scintillation tubes. After adding 3 mL of emulsifier-safe scintillation cocktail (Perkin Elmer), radioactivity was quantified with a TRI-CARB 2810TR (Perkin Elmer) scintillation counter.

The cholesterol secreted by astrocytes was assessed by measuring cholesterol contents in ACM by HPTLC and fluorimetry. For HPTLC, 1 mL of ACM was loaded and run. The bands were visualized and quantified using ChemidocTM Gel Imaging System and a standard concentration curve of cholesterol. The fluorimetric analysis was performed with the Amplex® Red cholesterol Kit (Thermo, catalogue # A12216). The assay is based on an enzyme-coupled reaction that detects free cholesterol and cholesteryl esters. Samples diluted in 1X reaction buffer were incubated with 50 μL of Amplex Red reagent working solution for 30 minutes at 37°C in the dark. The fluorescence was measured in a microplate reader Varioskan Lux. Excitation range was 530-560 nm and emission detection 590 nm The samples were assayed in triplicate. Filipin III was purchased from Sigma and used as per the manufacturer instructions.

### Statistical analysis

Each experiment was performed with cells from at least three independent cultures. Unpaired student’s T-test was used to compare one variable between groups, and two-way ANOVA followed by the multiple comparisons Tukey’s *post-hoc* test to compare two or three variables among groups. The interval of confidence was 95%, and data were represented as the mean ± standard error of the mean (SEM). A p-value < 0.05 was considered statistically significant; statistical differences are indicated as *p < 0.05, **p < 0.01, *** p < 0.001 and **** p < 0.0001.

## Results

### Dysregulation of neurodevelopmental pathways in CCALD and CAMN brain transcriptomes

We examined whether genes related to neurodevelopment, synaptic plasticity and neurotransmission are dysregulated in cortical transcriptomes from patients who died of CCALD or CAMN and age-matched controls^38^. To this end, the complete lists of differentially expressed genes (DEG) (p-value < 0.05) of CCALD or CAMN *vs* the respective controls (Suppl. data 1) were subjected to pathway (Reactome, KEGG) and ontology (GO-CC and GO-BP) analyses using the databases and search tools provided by Enrichrl^46^.

Several categories related to development, axonal growth, synaptic signaling and synaptic compartments were significantly dysregulated in CCALD (Fig. 1a, Suppl. data 4.) Some categories ranked among the first one of their groups; for example, ‘Developmental Biology’ was in the third position in Reactome, ‘Neuroactive ligand-receptor interaction’ was second in KEGG, and ‘Dendrite’ and ‘Neuron projection’ were, respectively, third and fourth in GO-Cellular Compartment. In addition, dysregulated categories related to ‘Programed cell death’ and ‘Apoptosis’ were added to the list of pathways potentially related to abnormal neural-circuit organization in X-ALD for two reasons. First, stochastic or lineage-determined cell death occurring at different stages of nervous-system development are essential for proper circuit configuration^47^. Second, VLCFA is toxic to brain cells *in vitro*^4–8^, suggesting that VLCFA may exacerbate programmed brain cell death *in vivo*. The pool of DEG related to neurodevelopment represents 6.0% of the DEG in CCALD (Fig. 1a, volcano plot, points in red).

**Figure 1.**
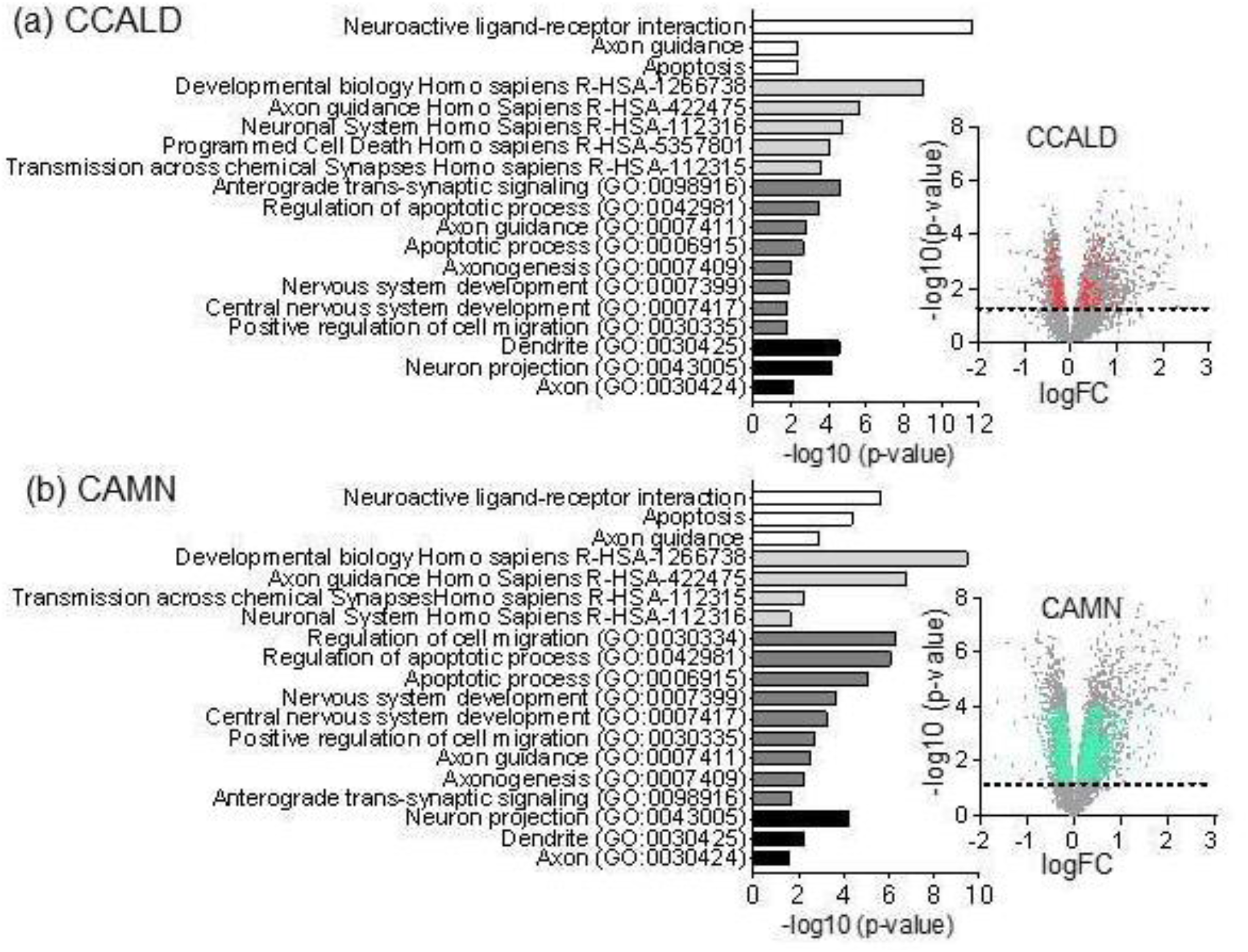
Neurodevelopment-centered pathway and ontology analyses of CCALD and CAMN transcriptomes. Dysregulated categories related to neurodevelopment and neurotransmission in transcriptomes from CCALD (a) and CAMN (b) patients vs age-matched controls using DEG (p < 0.05.) Categories from KEEG, Reactome Homo Sapiens (R-HAS), GO-Biological Property, and GO-Cellular Compartment are ranked according to the −log10 of p-values and labeled in different shades of grey. Volcano plots (right) show in red (CCALD) and blue (CAMN) the distribution of DEG related to neurodevelopment and neurotransmission in the total transcriptome. Dashed lines indicate genes p < 0.05. The pool of DEG related to neurodevelopment and neurotransmission represent 6% of DEG in CCALD and 22.7% in CAMN.

Most of the development categories dysregulated in CCALD were also dysregulated in CAMN and ranked among the first 10 categories altered in their respective groups (Fig. 1b, Suppl. data 5.) The pool of DEG related to neurodevelopment represents 22.7% of the DEG in AMN (Fig. 1b, volcano plot, points in blue). All in all, the data from patients with X-ALD point to the existence of congenital defects in neural circuits that persist into adulthood.

### Depletion of Abcd1/2 in neurons and survival assessment

To validate the predictions from the transcriptomic data, we examined the direct impact of *Abcd* deficiency on neuron survival and development in cultures of neonatal rat cortical neurons devoid of microglia and oligodendrocytes and with low astrocyte abundance (See Methods). The *in vitro* model of X-ALD-like neurons was generated by silencing *Abcd1 and Abcd2* transporters by transduction with viral vectors of shRNAs and/or exposure to 50 μM C26:0 to achieve the typically high concentrations of VLCFA present in X-ALD^50^ (Fig. 2a, c.) *Abcd1* and *Abcd2* transporters were silenced in unison. The influence of astrocyte-secreted factors was mimicked by exposing neurons to ACM, either from control astrocytes or from the henceforth called X-ALD-like astrocytes, which fulfilled two conditions: their *Abcd1/2* transporters were silenced with transfected siRNAs and they were exposed to a pulse of 50 μM C26:0 (Fig. 2b, c.) Silencing effectiveness was assessed in all cases after the assays were concluded (Fig. 2c.)

**Figure 2.**
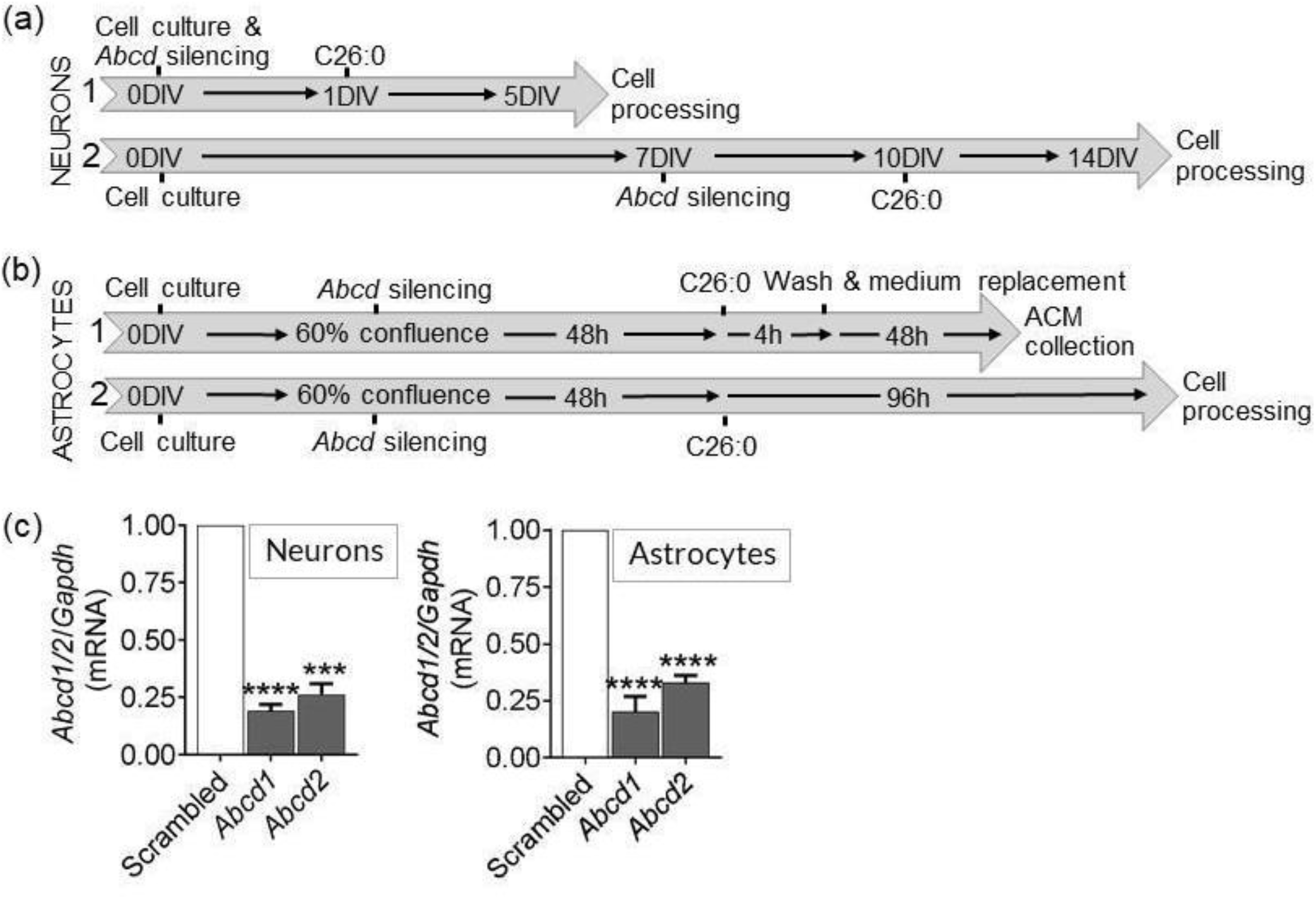
*In vitro* models to study neuronal development in X-ALD. (a) *Experimental outline for neurons.* Neuronal cultures were generated from neonatal rat cortices. The expression of *Abcd1* and *Abcd2* transporters was silenced with shRNAs transduced by adeno-associated viral vectors. Neurons were infected with the viral vectors at 0 days-in-division (DIV) for short-term studies (e.g., dendrite arborization and spinogenesis) (a.1) and at 7DIV for long-term ones (e.g., axonogenesis) (a.2.) The X-ALD-like phenotype was enhanced by exposure to 50 μM C26:0 dissolved in α-cyclodextrin (‘α-cyclo’) as a vehicle for four days before cell processing. Control cells were treated with scrambled shRNA and vehicle a-cyclo. The effect of astrocytes on neurons was examined using astrocyte conditioned media (ACM) from control or *Abcd1/2* silenced astrocytes exposed to a 4-hour pulse of vehicle (ACM^CRTL^) or C26:0 (ACM^X-ALD^) (see b.) (b) *Experimental outline for astrocytes.* Astrocytic cultures were generated from neonatal rat cortices. The expression of *Abcd1* and *Abcd2* transporters was silenced with transfected siRNA. The X-ALD-like phenotype was enhanced by exposure to 50 μM C26:0 dissolved in the vehicle α-cyclodextrin (‘α-cyclo’.) Control cells were treated with scrambled siRNA and vehicle. Cells were subjected to two protocols (1, 2) either to collect astrocyte conditioned media (ACM) or to study astrocytes themselves. In the first case C26:0 was washed out and ACM were collected for another 48 h. (c) *Efficient Abcd1 and Abcd2 co-silencing in neurons and astrocytes.* The expression of *Abcd1* and *Abcd2* was silenced using shRNA (neurons) and siRNA (astrocytes) for each transporter. *Abcd1* and *Abcd2* expression was tested with qPCR. Data are the means ± SEM, 3-4 independent experiments. *** p < 0.001 and **** p < 0.001, Unpaired T-test.

The dysregulation of the category ‘apoptosis’ in the CCALD and AMN arrays (Fig. 1a, b) prompted us to first examine whether *Abcd1/2* deficiency increased the death rate of neurons and astrocytes. Apoptotic cellular death was measured with TUNEL staining.

Basal neuronal death in the neuronal cultures was around 15%, α-cyclo caused an additional 10% of neuronal death, and the infections with viral vectors another 25%. That is, both the vehicle and the infections increased neuronal death over basal conditions up to 50% in the group shScrambled + α-cyclo, in the absence of ACM. This was considered the control group to test the effects of *Abcd1/2* silencing, C26:0 and ACM. Neuronal death increased by 28% (p < 0.05) upon addition of C26:0, and by 90% (p < 0.0001) upon silencing of *Abcd*1/2 transporters (Fig. 3a, b.) Addition of C26:0 in the latter group did not increase neuronal death. Presence of ACM^CRTL^ decreased cellular death in shScrambled treated with C26:0 and in all silenced conditions. For example, the increased death of *Abcd1/2* deficient neurons was reduced by 66% in the presence of ACM^CRTL^ (p < 0.0001.) There was no statistically significant difference between ACM^CTRL^ and ACM^X-ALD^—although the latter was slightly less protective, as suggested by the larger p-values associated (p-values of all post-hoc analyses are in Suppl. data 6.)

**Figure 3.**
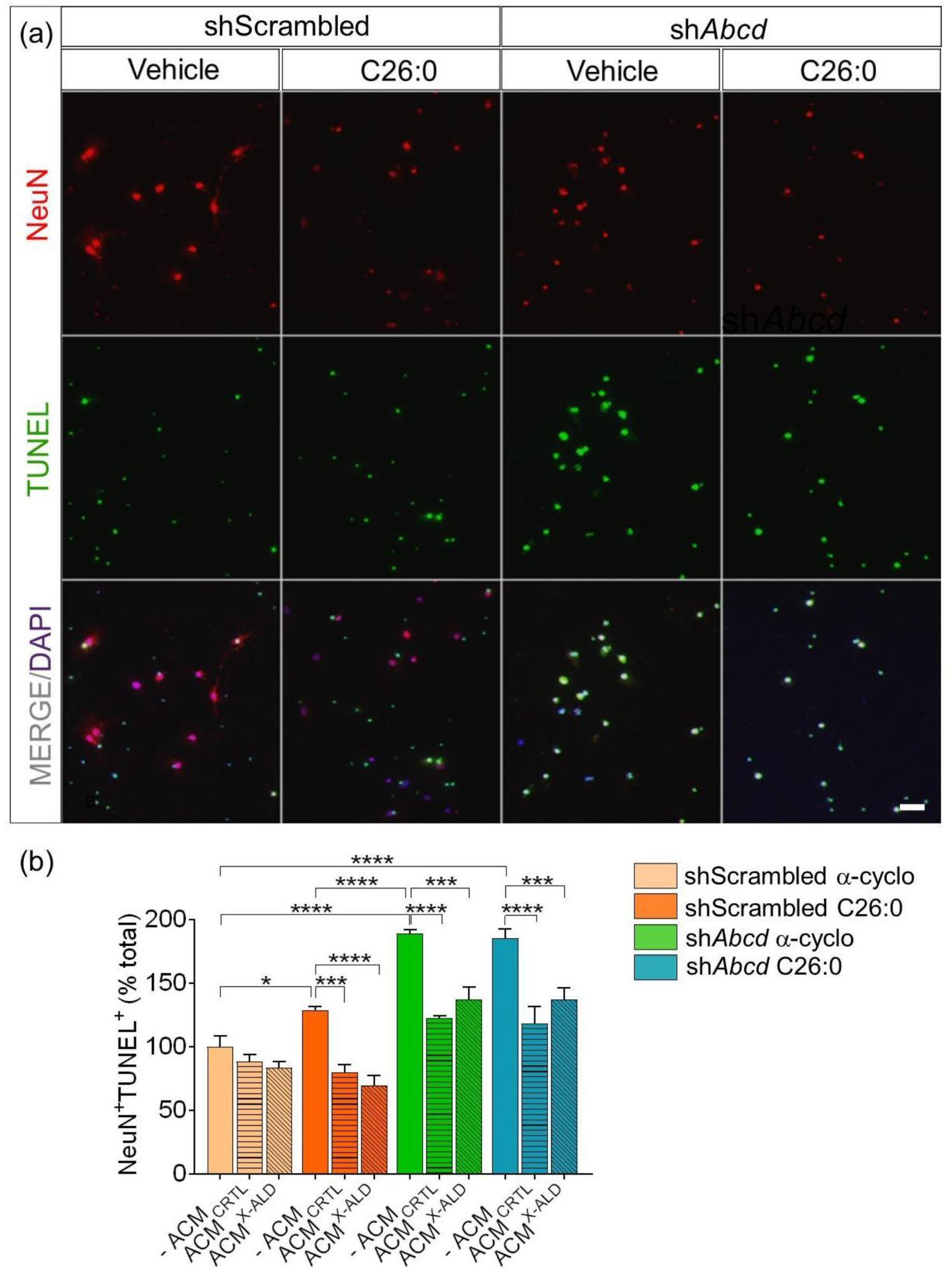
Increased death rate in *Abcd1/2* deficient neurons. Neuronal death was assessed by co-localization of TUNEL (green) and NeuN-positive cells in four experimental conditions as indicated by colors. Data are expressed as percentage of ShScrambled + α-cyclo values in the absence of ACM. Warm colors (salmon/orange) are neurons transduced with shScrambled, and cold colors (green/blue) neurons transduced with *shAbcd1/2.* Each condition was exposed to either α-cyclo or C26:0. The impact of astrocytes was tested with astrocyte-conditioned media (ACM), either from control (ACM^CRTL^) or *Abcd1/2* deficient astrocytes (ACM^X-ALD^). (a) Representative images of neuronal death. Scale bar is 25 μm. Notice the increase in TUNEL positive cells in the *shAbcd1/2* conditions. (b) Quantitation of neuronal death. A minimum of 300 cells was analyzed per experiment. Data are the means ± SEM, n = 3 independent cultures. Comparisons among groups were performed with two-way ANOVA followed by post-hoc Tukey’s multiple comparisons; all p-values are detailed in Suppl. data 5, while, for clarity, only statistics for three types of comparisons are presented in the figure: effect of C26:0 (siScrambled / *minus* ACM/vehicle *vs* siScrambled / *minus* ACM / C26:0), showing slight exacerbation of neuronal death in the presence of the VLCFA; effect of *Abcd1/2* deficiency (sh*Abcd vs* shScrambled in *minus* ACM conditions) showing pronounced increases in neuronal death; and protective effects of both ACM^CRTL^ and ACM^X-ALD^ versus *minus* ACM, both when neurons are exposed to C26:0 or are *Abcd1/2* deficient; * p < 0.05; ** p < 0.01; *** p < 0.001; **** p < 0.0001.

By contrast, extreme X-ALD-like conditions (deletion of *Abcd1/2* transporters *plus* exposure to high concentrations of C26:0) did not increase astrocyte death over the low basal death rate of 0.5% (Fig. 4a, b.)

**Figure 4.**
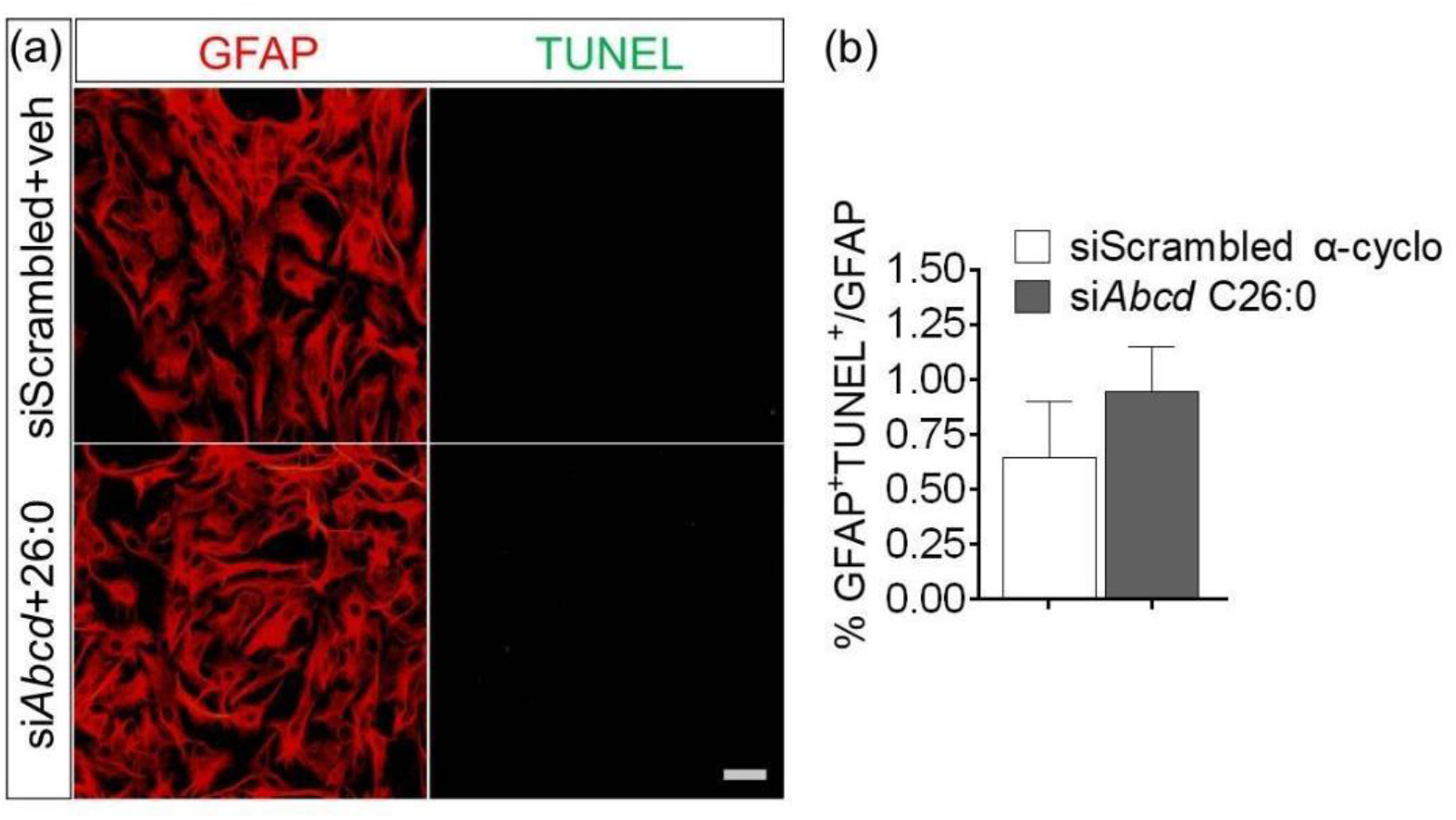
No increased death rate in *Abcd* deficient astrocytes. Astrocyte death was assessed by co-localization of TUNEL (green) with GFAP (red) in two conditions: siScrambled + α-cyclo and *siAbcd1/2* + 50 μM C26:0. (a) Representative images. Scale bar is 25 μm. (b) Quantification of astrocyte death. A minimum of 300 astrocytes was counted *per* independent experiment. Data are the means ± SEM, n = 3 independent cultures, Unpaired T-test. No TUNEL+ cells were detected over basal rate.

So far, we conclude: (i) *Abcd1/2* deficiency increases neuronal death in cultures, i.e., there is a neuron-autonomous pathological condition associated with loss of *Abcd1/2* function; (ii) the predominant action of ACM with regards to neuronal demise is protection, thus ruling out the possibility that astrocytes may secrete (or not sufficiently metabolize) factors that are toxic to neurons in X-ALD, as shown in experimental models of ALS^48^ and septic shock^49^; and (iii) C26:0 mildly exacerbated the neuronal death resulting from *Abcd1/2* depletion, but does not induce astrocyte death, at variance with a previous report^6^.

### Morphometric analysis of neuronal development upon Abcd1/2 silencing

Because aberrant neurodevelopment can manifest as abnormal neurite generation in addition to neuronal death, we analyzed dendrite arborization and diameter, spinogenesis, and axonal growth in *Abcd1/2* deficient neurons ± C26:0. We also compared the effects of ACM^CTRL^ *vs* ACM^XALD^ to distinguish neuron-autonomous from astrocyte-dependent alterations, for neuritogenesis may be a more sensitive readout than cell death to unravel subtle differences between ACM^CTRL^ *vs* ACM^X ALD^.

#### Effect of Abcd1/2 deficiency

Arborization was assessed with Sholl analysis, based on the number of intersections of dendrites with concentric circles placed every micrometer from the neuronal center of mass to the farthest point of the longest neurite (Fig. 5b.) Control ShScrambled neurons had 12 ramifications and 450-500-μm long dendrites, whereas Sh*Abcd* neurons had up to 10 ramifications and 300-400 μm long dendrites (Fig. 5c.) Note the displacement of green/blue curves to the left as compared to salmon/orange curves in Fig. 5c. Post hoc analyses confirmed statistically significant differences between shScrambled vs *shAbcd* neurons in all paired comparisons: ShScrambled/α-cyclo vs ShAbcd/α-cyclo; ShScrambled/C26:0 vs ShAbcd/C26:0 and ShScrambled/α-cyclo vs ShAbcd/C26:0 (Fig. 5d-f without SEM for visual clarity.)

**Figure 5.**
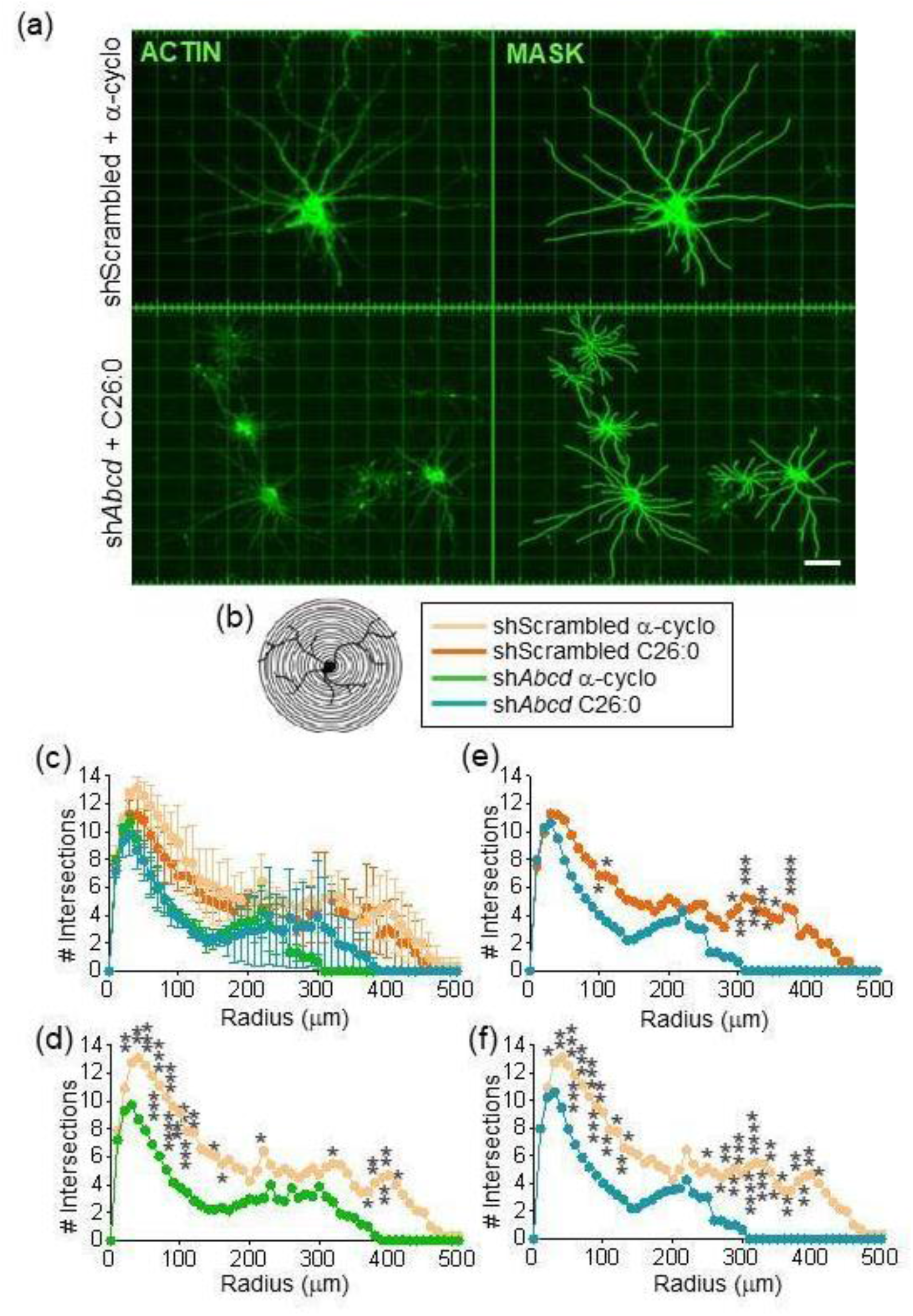
Defective dendrite arborization in *Abcd1/2* deficient neurons. Neurons were infected with shScrambled- or *shAbcd*-harboring viral vectors and exposed at 14 DIV to α-cyclodextrin, ‘α-cyclo’ or 50 μM C26:0. Data of neurons grown in ACM^CTRL^ are shown. ACM^X-ALD^ did not significantly affect the defective arborization elicited by *Abcd* deficiency with respect to ACM^CTRL^ (data not shown.) (a) Representative images of neurons labeled with ActinGreen™488 and visualized using confocal microscopy. Morphometric analyses were performed using the ‘filament tracer’ plug-in of IMARIS, which builds masks out of fluorescence. The grid is the tool used by IMARIS to measure relative distances. Scale bar is 25 μm. *Abcd* deficient neurons occupied smaller surfaces than control ones. (b) Sholl graph. Intersections of neurites with circles were counted as a readout of arborization. (c) Quantification of dendrite arborization as the number of dendrite intersections with Sholl circles every 1 μm from the center of the neuronal soma. Data are the means ± SEM of n=3 independent experiments, each experiment being the mean and SEM of 15 neurons. Warm colors are shScrambled neurons and cold colors sh*Abcd* neurons (d, f) Paired comparisons without SEM for visual clarity. (d) shows shScrambled + α-cyclo *vs* sh*Abcd* + α-cyclo; (e) shScrambled + C26:0 vs sh*Abcd* + C26:0; and (f) shows the comparison of extreme conditions shScrambled + α-cyclo vs sh*Abcd* +C26:0. * p < 0.05, **p < 0.01, *** p < 0.001 and **** p < 0.0001 (Two-way ANOVA and post-hoc Tukey’s multiple comparisons test.) The distribution of intersections in *Abcd*-deficient neurons shifted to the left as compared to shScrambled neurons in all comparisons, indicating fewer ramifications in silenced neurons. Addition of C26:0 shortened neurites in *Abcd*-silenced conditions (max length is 400 μm in the green curve and 300 μm in the blue one.)

The plug-in ‘filament tracer’ of IMARIS marks the dendrite shaft and its protrusions (i.e., spines) with different colors, thus allowing independent assessments of shaft diameter and spine density (Fig. 6a.) *Abcd* deficiency reduced shaft diameters. In all paired comparisons, 80% of the dendrites of shScrambled neurons, but only 40-60% of *Abcd* silenced neurons, were thicker than the minimum diameter that can be detected, 0.35 μM. Note the shift to the left of the green/blue curves (Fig. 6b-e.)

**Figure 6.**
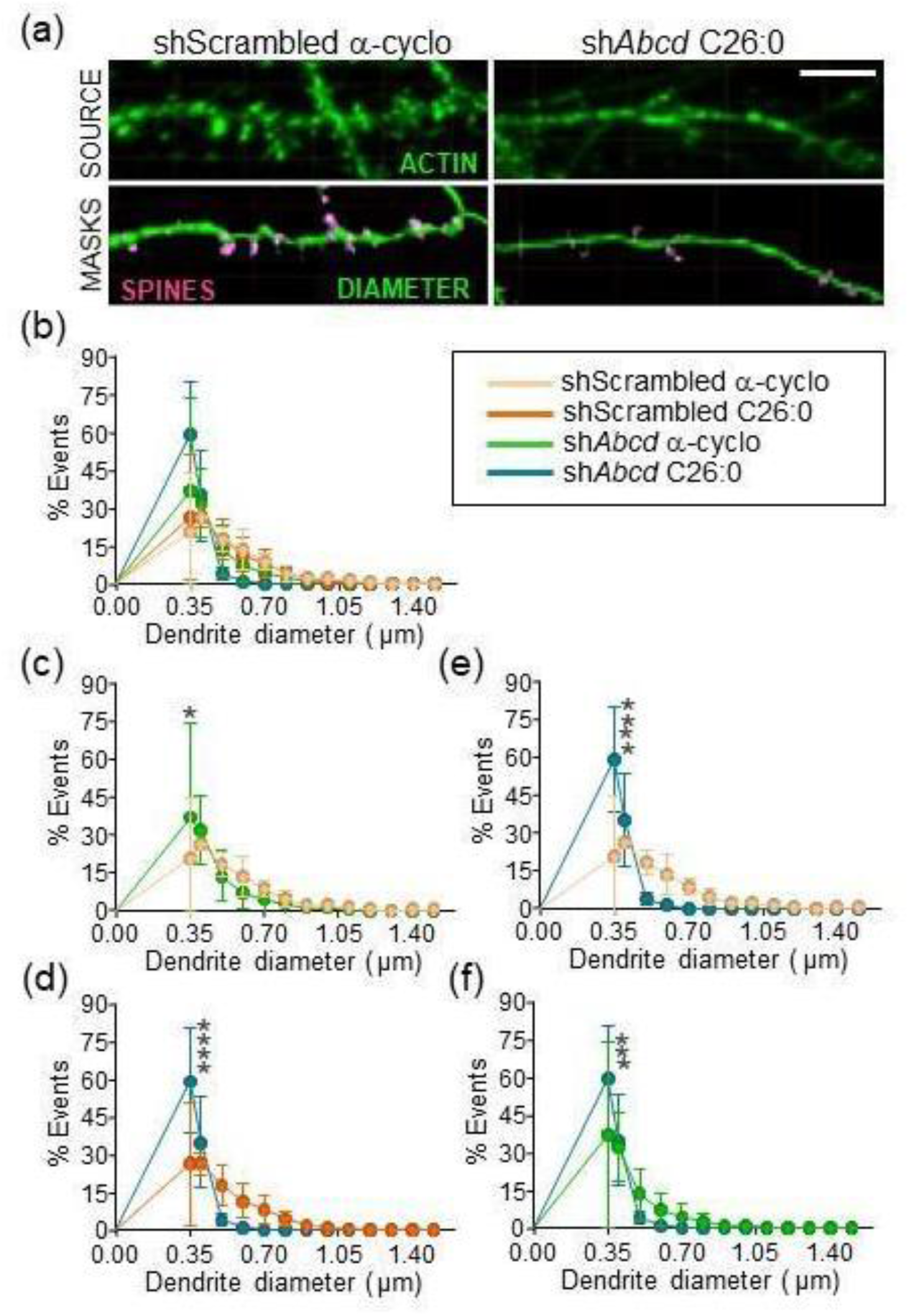
Reduced dendrite diameter in *Abcd1/2* deficient neurons. (a) Top, representative images of neuronal dendrite in extreme experimental conditions (shScrambled + α-cyclo vs sh*Abcd* + C26:0) immunostained for actin (green). Bottom, masks of shafts (green) and spines (pink). Scale bar is 5 μm. (b) Quantification of shaft diameters showing all experimental groups. Data of cells grown in ACM^CRTL^ are shown (for this readout, data in the presence of ACM^X-ALD^ were not significantly different to those obtained with ACM^CRTL^.) Data are the means ± SEM of n=3 independent experiments, each experiment is the mean and SEM of 15 neurons. Warm colors are shScrambled neurons and cold colors *shAbcd1/2* neurons (c, f) Paired comparisons and post hoc statistics without SEM for visual clarity. *Abcd1/2* silencing significantly diminished shaft diameter both in the presence of vehicle (c) or C26:00 (d) or when comparing extreme conditions (e). C26:0 exacerbated the effect of silencing as deduced from the significant reduction of diameter when comparing silenced neurons treated with a-cyclo with silenced neurons treated with C26:0 (f). * p < 0.05, **p < 0.01, *** p < 0.001 and **** p < 0.0001 (Two-way ANOVA and post-hoc Tukey’s multiple comparisons test.)

We quantified spinogenesis as the distribution of the probability of events in dendritic arbors (Fig. 7.) One event is the percentage of dendrites with a given density of spines (spines per μm of shaft.) Densities were divided in discrete intervals (0 - 0.20, 0.21 - 0.40, 0.41 - 0.60, and so on, up to 2.8 - 3.0.) For example, 20 events in the 0-0.20 range means that 20% of the dendrites had 0-0.20 spines/μm. Most of the dendrites fell in the 0.2-1.2 range. ShScrambled neurons (salmon/orange) peaked around 0.8 and *shAbcd* neurons (green/blue) around 0.5; in other words, 80% of control neurons had 0.4-1.2 spines/μm, with 40% presenting spine densities greater than 0.8, whereas 80% of *Abcd* deficient neurons had 0.2-1 spines/μm, with 50% of this population showing 0.2-0.6 spines/μm (Fig. 7c-e.) This shift indicates reduced density of spines upon *Abcd1/2* silencing. Post hoc comparisons between shScrambled and sh*Abcd* neurons are shown in Figs.7c-h.

**Figure 7.**
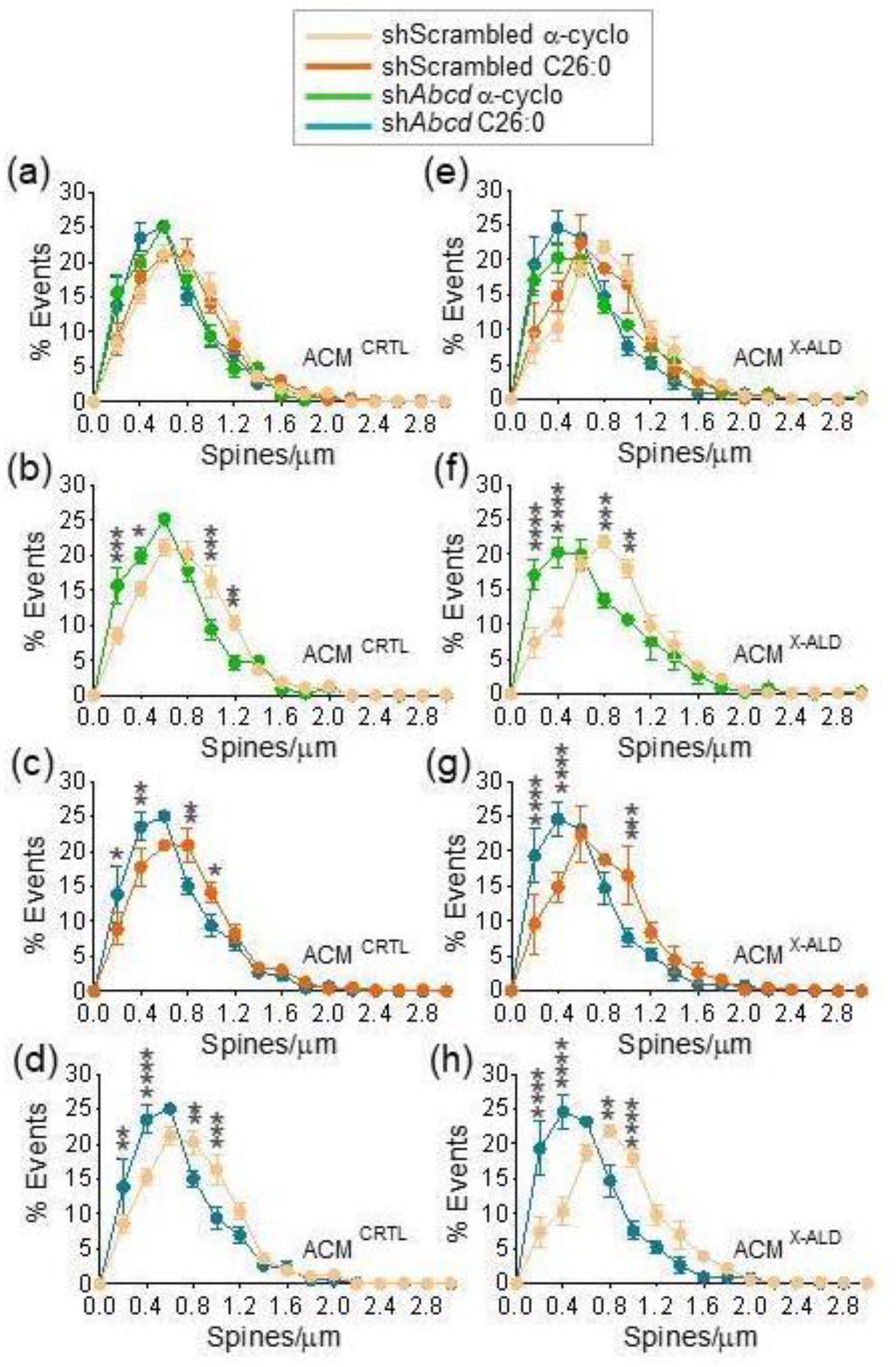
Reduced spinogenesis in *Abcd1/2* deficient neurons. X axis are the intervals of number of spines per μm and Y are the percentage of dendrites (events) presenting the density of spines at a given interval. (a, e) Distribution of spine densities in all experimental conditions in the presence of ACM^CRTL^ (a) or ACM^X-ALD^ (e). (b-h) Individual comparisons of (a) or (b) showing post hoc statistics (Two-way ANOVA followed by Tukey). The distribution of events in *Abcd* null neurons shifted to the left, indicating reduced spine numbers per μm and thinner dendrites. The greater significance of some points in e as compared to d reveals a mild but detectable reducing impact of C26:0 on spine density. ACM^X-ALD^ mildly exacerbated the effect of *Abcd* silencing, as shown by the deepest separation of ShScrambled and Sh*Abcd* curves in ACM^X-ALD^ with respect to ACM^CRTL^. Likewise, the addition of C26:0 slightly worsened the effect of *Abcd* deficiency (d vs b).

Finally, axonal length was 30% shorter in X-ALD-like neurons (*Abcd1/2* silenced + C26:0 in ACM^X-ALD^) than in controls (shScrambled + α-cyclo in ACM^CRTL^) (p-value 0.0001, Fig. 8.) Only extreme X-ALD-like conditions and not intermediate ones (e.g., *Abdc* silencing but not C26:0) were compared with controls in the variable ‘axonogenesis’ because axon growth requires long-term cultures that are difficult to maintain due to the limited viability of neuronal cultures.

**Figure 8.**
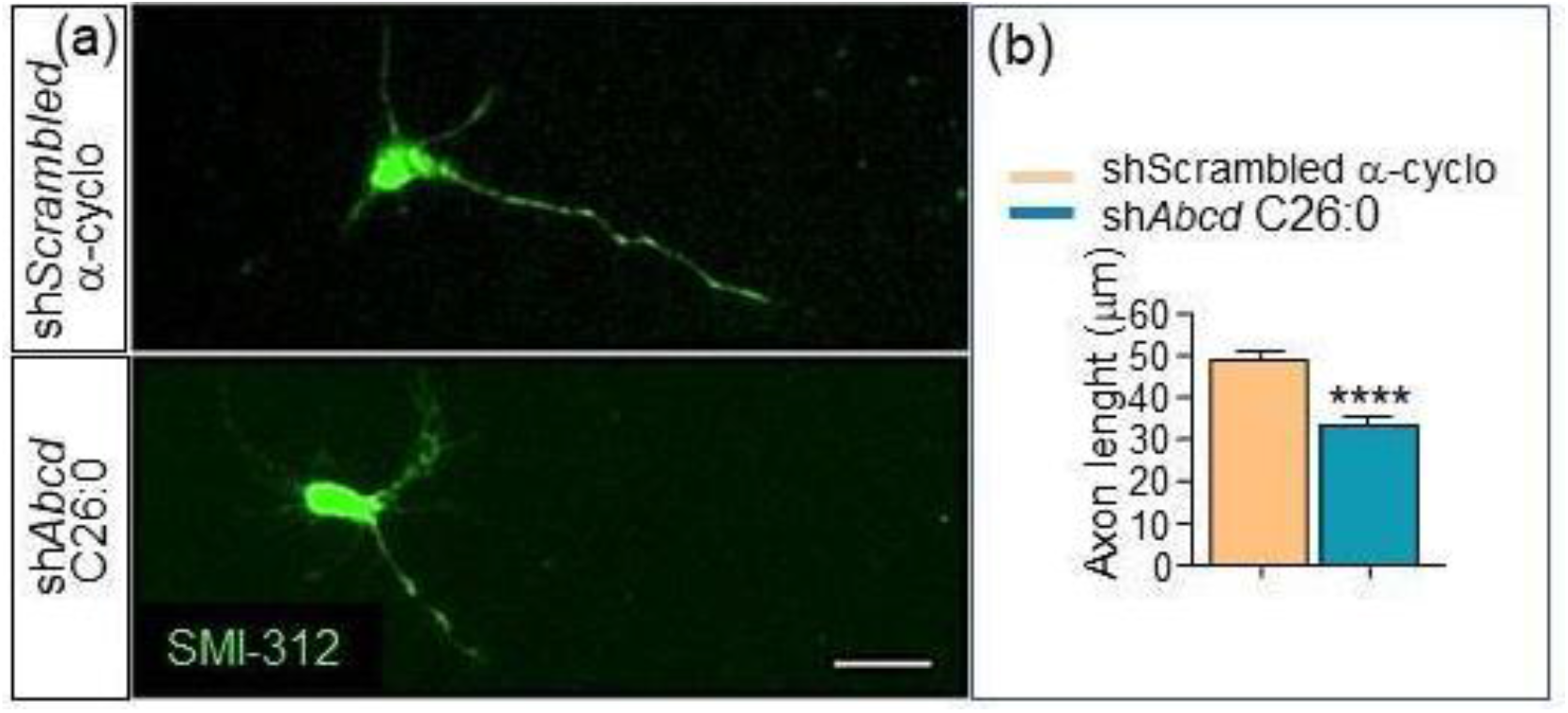
Reduced axonal growth in *Abcd1/2* deficient neurons. Representative images (a) or neurons stained with SMI-312 and fluorescence quantification (b) of axonal length. X-ALD neurons are shAbcd + C26:0 and controls are shScrambled + α-cyclo. Scale bar 10 μm. ****p<0.0001, Unpaired T-test. N=3, 200-300 neurons measured per experiment.

#### Effect of C26:0

In dendrite arborization, C26:0 shortened the length of the longest neurite in *Abcd* silenced conditions from 400 to 300 μm (Fig. 5, green vs blue curves). In dendritic shafts, the lowest percentage of silenced neurons with dendrites thicker than 0.35-μm (40%) was observed in the presence of C26:0 (blue line in Fig. 6d-f), suggesting that the VLCFA exacerbated the effect of *Abcd* silencing. Likewise, C26:0 slightly worsened the effect of *Abcd* silencing on spinogenesis, as shown by the greater statistical significance in some intervals when comparing extreme conditions (shScrambled + α-cyclo vs *shAbcd1/2* + C26:0) (Fig.7d) than when comparing paired conditions α-cyclo/α-cyclo (Fig. 7b) and C26:0/C26:0 (Fig. 7c). For example, at a density of 0.5 spines/μm, the p-value of the effect caused by *Abcd* silencing is p< 0.05 in the presence of α-cyclo (Fig. 7b), p<0.01 in the presence of C26:0 (Fig. 7c), and p<0.0001 when control neurons were treated with α-cyclo and sh*Abcd* neurons with C26:0 (Fig. 7d).

#### Effect of ACM

No difference between ACM^CRTL^ and ACM^X-ALD^ was detected in dendrite thickness and arborization, but ACM^X-ALD^ mildly exacerbated the effect *Abcd* silencing on spinogenesis, as the shift to the left of green and blue curves is more pronounced in Fig. 7 e-h than in a-d.

In summary, as per morphological criteria, *Abcd* deficiency globally impairs neuronal development, while ACM^X-ALD^ slightly worsens spinogenesis. If one extends these observations to the real human disease, in which *ABCD* deficient neurons would be under the influence of *ABCD* deficient astrocytes, the data pinpoint the existence of two elements altering the proper morphological development of neurons: a most prominent one, intrinsic to neurons, slightly exacerbated by excess of VLCFA, and a lesser astrocytic component specifically affecting spine formation.

### Screen of pathways involved in aberrant neurodevelopment

Next, we aimed to gain insight into signaling pathways accounting for the neuron-autonomous defective neuritogenesis and impaired survival. Again, we looked up in the CCALD database for clues, for brains from children are still developing. We focused on genes related to the main signaling pathways governing nervous system development at several stages and anatomical levels, including neuronal maturation, morphogenesis of dendritic arbors, axon guidance and synaptogenesis: *Wnt*^51^, Notch^52^, Sonic hedgehog (*Shh*)^53^, neurotrophins^53^, *Stat3*^54,55^.

Several genes related to these pathways were differentially expressed (p < 0.05) in CCALD (Table 1, organized in alphabetical order.) *Ephb2, Notch3, Ntrk2, Shh, Wnt2* and *Wnt6* were downregulated and Ntrk3 and *Stat 3* upregulated. Several of these genes (*Ephb2, Notch3, Ntrk2, Stat3* and *Wnt6*) were also dysregulated in CAMN (Table 1), suggesting that pathways formerly involved in development, that in the adult brain specialize in regulating synaptic plasticity, learning and memory and repair^56^, are altered in adult carriers of mutated *ABCD* genes.

**TABLE 1.**
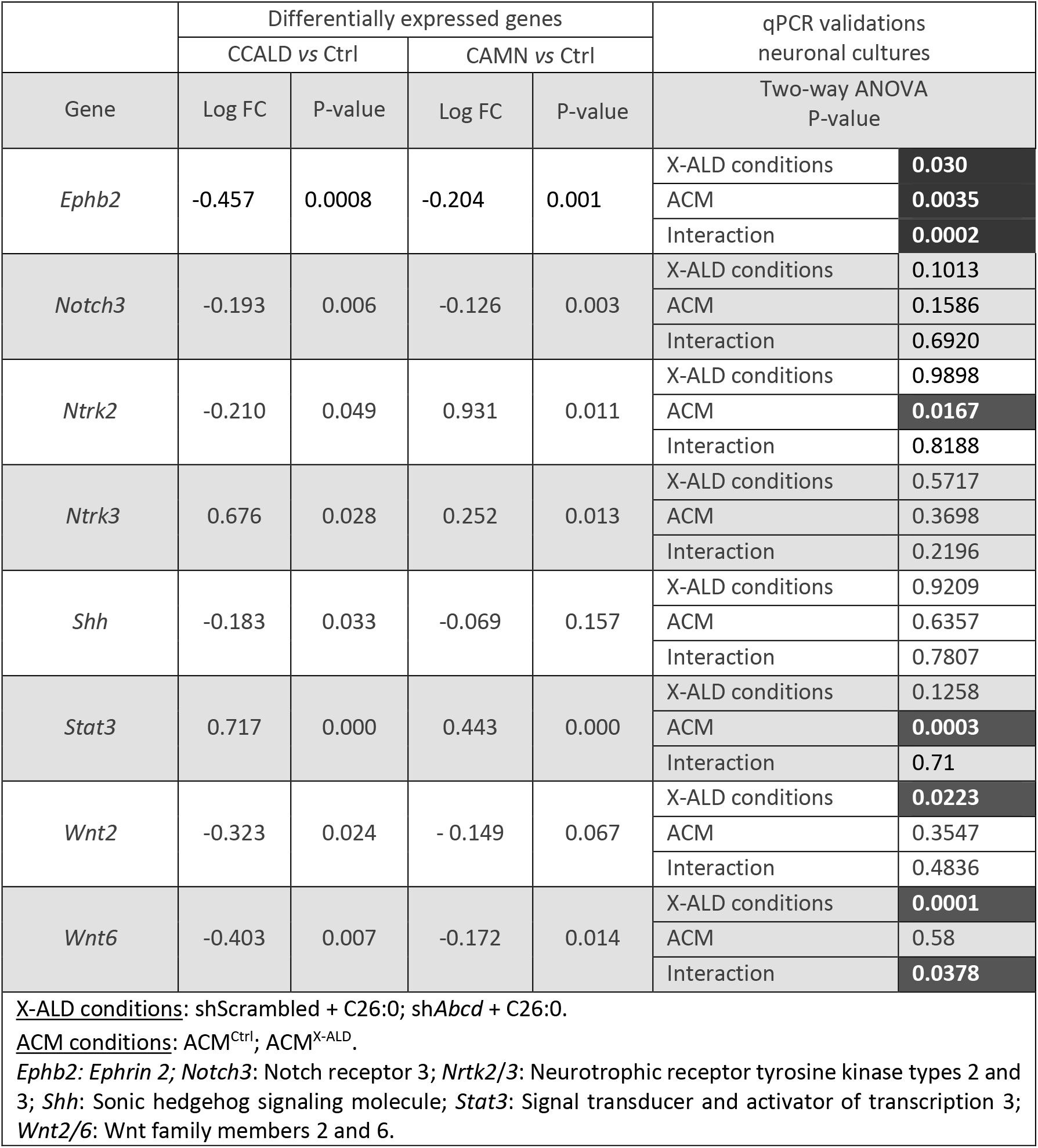
Expression of genes related to CNS development.

| TABLE 1. Expression of genes related to CNS development |  |  |  |  |  |  |
| --- | --- | --- | --- | --- | --- | --- |
|  | Differentially expressed genes |  |  |  | qPCR validations<br>neuronal cultures |  |
|  | CCALD vs Ctrl |  | CAMN vs Ctrl |  |  |  |
| Gene | Log FC | P-value | Log FC | P-value | Two-way ANOVA<br>P-value |  |
| *Ephb2* | -0.457 | 0.0008 | -0.204 | 0.001 | X-ALD conditions | **0.030** |
|  |  |  |  |  | ACM | **0.0035** |
|  |  |  |  |  | Interaction | **0.0002** |
| *Notch3* | -0.193 | 0.006 | -0.126 | 0.003 | X-ALD conditions | 0.1013 |
|  |  |  |  |  | ACM | 0.1586 |
|  |  |  |  |  | Interaction | 0.6920 |
| *Ntrk2* | -0.210 | 0.049 | 0.931 | 0.011 | X-ALD conditions | 0.9898 |
|  |  |  |  |  | ACM | **0.0167** |
|  |  |  |  |  | Interaction | 0.8188 |
| *Ntrk3* | 0.676 | 0.028 | 0.252 | 0.013 | X-ALD conditions | 0.5717 |
|  |  |  |  |  | ACM | 0.3698 |
|  |  |  |  |  | Interaction | 0.2196 |
| *Shh* | -0.183 | 0.033 | -0.069 | 0.157 | X-ALD conditions | 0.9209 |
|  |  |  |  |  | ACM | 0.6357 |
|  |  |  |  |  | Interaction | 0.7807 |
| *Stat3* | 0.717 | 0.000 | 0.443 | 0.000 | X-ALD conditions | 0.1258 |
|  |  |  |  |  | ACM | **0.0003** |
|  |  |  |  |  | Interaction | 0.71 |
| *Wnt2* | -0.323 | 0.024 | - 0.149 | 0.067 | X-ALD conditions | **0.0223** |
|  |  |  |  |  | ACM | 0.3547 |
|  |  |  |  |  | Interaction | 0.4836 |
| *Wnt6* | -0.403 | 0.007 | -0.172 | 0.014 | X-ALD conditions | **0.0001** |
|  |  |  |  |  | ACM | 0.58 |
|  |  |  |  |  | Interaction | **0.0378** |
| X-ALD conditions: shScrambled + C26:0; shAbcd + C26:0. |  |  |  |  |  |  |
| ACM conditions: ACM<sup>Ctrl</sup>; ACM<sup>X-ALD</sup>. |  |  |  |  |  |  |
| *Ephb2*: Ephrin 2; *Notch3*: Notch receptor 3; *Ntrk2/3*: Neurotrophic receptor tyrosine kinase types 2 and 3; *Shh*: Sonic hedgehog signaling molecule; *Stat3*: Signal transducer and activator of transcription 3; *Wnt2/6*: Wnt family members 2 and 6. |  |  |  |  |  |  |

We validated the DEG from CCALD and dissected out the neuronal vs astrocytic influence in our *in vitro* model of neuronal development by qPCR (Fig. 9a). *Syn* was assayed alongside as a positive control of reduced synaptogenesis, because it encodes synaptophysin, a synaptic vesicle component commonly used as a marker of synapse maturation (Fig. 9b.) Two-way ANOVA was used to determine the effects of ‘X-ALD-like conditions’ (shScrambled α-cyclo vs *shAbcd1/2* C26:0, for simplicity only extreme conditions were only used herein) and ‘ACM’ (ACM^CTRL^ *vs* ACM^XALD^) in the qPCR-based validations.

**Figure 9.**
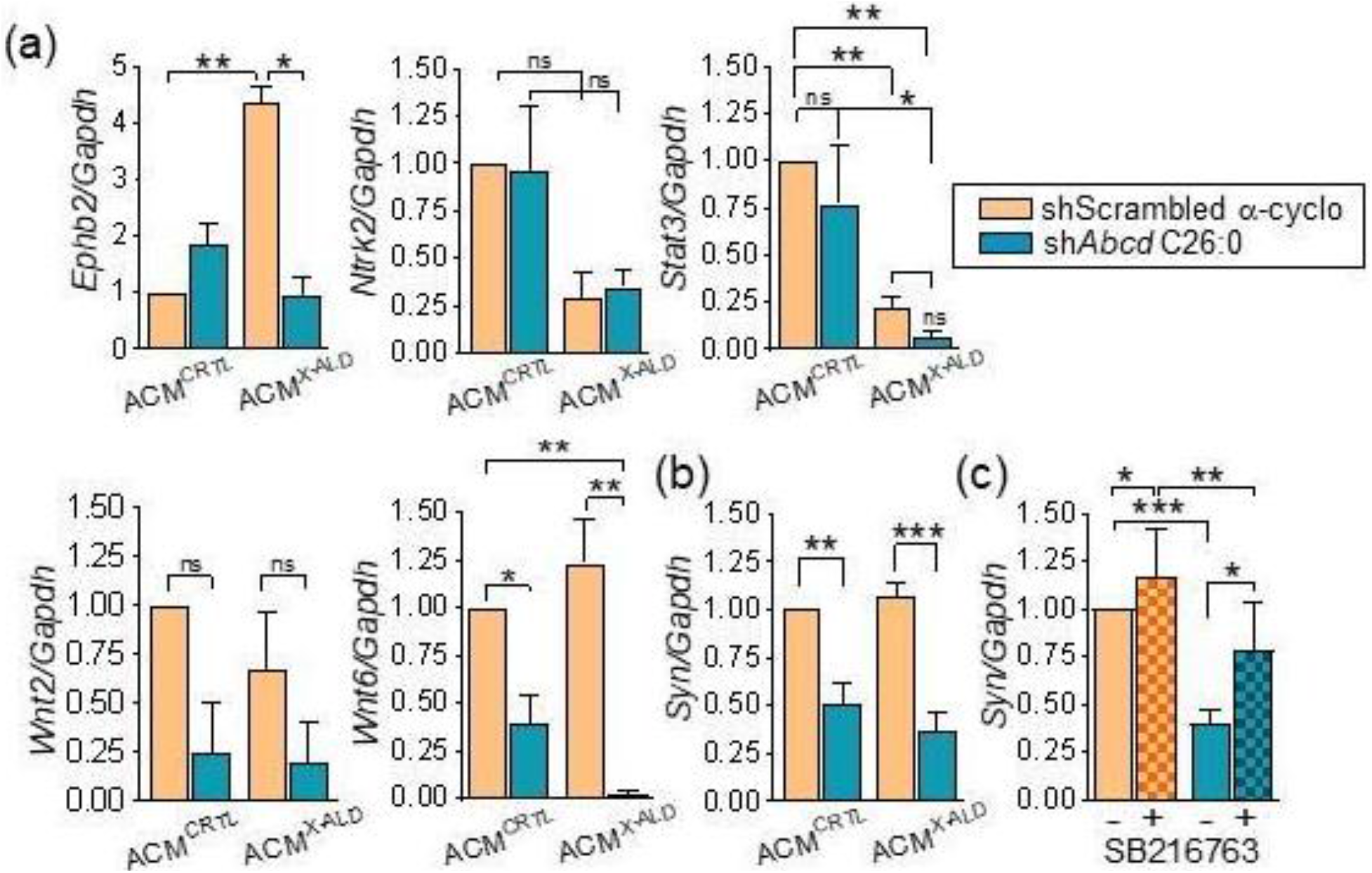
Validation of neurodevelopmental DEG in CCALD in *Abcd1/2* deficient neurons. a) Quantification of mRNA contents of DEG from Table 1 in control (ShScrambled / α-cyclo) and X-ALD-like neurons (*shAbc1/2d* + C26:0) by qPCR in the presence of astrocyte conditioned media (ACM) from control (ACM^CTRL^) or si*Abdc1/2*/C26:0 astrocytes (ACM^X-ALD^.) *Gapdh* was used to normalize mRNA data. Data are the means ± SEM, n=3; data were analyzed with two-ANOVA (p-values of the impact of variables ‘X-ALD conditions’ and ‘ACM’ are in Table 1), followed by Tukey multiple comparisons test. Only genes that were statistically significant in one parameter of two-way ANOVA are shown. ‘X-ALD conditions’ had an impact on *Wnt2* and *Wnt6*, ‘ACM conditions’ on *Stat3* and *Nrtk2* and ‘interaction’ on *Ephb2* and *Wnt6;* *p < 0.05; **p < 0.01; ***p < 0.001. *Ephb2:* Ephrin 2; *Nrtk2:* Neurotrophic receptor tyrosine kinase type 2; *Stat3:* Signal transducer and activator of transcription 3; *Wnt2/6:* Wnt family members 2 and 6. b) *Syn*: Synaptophysin, showing reduction upon *Abcd1/2* silencing, which was more statistically significant in the presence of ACM^X-ALD^. c) Effect of the GSK3 inhibitor SB216763 on *Syn* expression in X-ALD neurons (ACM^X-ALD^). SB216763, used at 5 μM, was administered to neurons daily for three days before the RNA extraction. Data are the means ± SEM, n=3, two-way analysis followed by Tukey’s multiple comparisons; *p < 0.05; **p < 0.01; ***p < 0.001; ****p < 0.0001. The GSK3 inhibitor partially reversed the downregulation of *Syn* in X-ALD-like conditions and increased *Syn* expression in control neurons.

The variable ‘X-ALD-like conditions’ had an impact on the expression of *Wnt2* and *Wnt6*, ‘ACM’ impacted *Stat3* and *Ntrk3* expression, and there was a statistically significant interaction between the two variables in the case of Wnt6, and particularly *Ephb2* (Table 1). Post hoc comparisons of these genes are shown in Fig. 9a and follow three patterns. One, prominent up-regulation in ACM^X-ALD^, but only in ShScrambled neurons, shown by Ephb2. Two, downregulation in ACM^X-ALD^ in both shScrambled and *Abcd* silenced neurons, as predicted by two-way ANOVA, shown by *Stat3* and *Nrkt3*-although post hoc differences only were statistically significant in the case of *Stat3.* Three, down-regulation only in *Abcd* silencing with no or mild exacerbation in ACM^X ALD^, affecting Wnts. *Syn* followed this latter pattern, too (Fig. 9b.) The downregulation caused by *Abcd* deficiency was slightly more pronounced in ACM^X ALD^ (p<0.0004) than in ACM^CRTL^ (p<0.0021), consistent with the effect of ACM^X ALD^ on spinogenesis described above.

Wnt proteins are secreted lipid-modified signaling glycoproteins that act by activation of Frizzled receptors in cellular membranes. Frizzled activation frees β-catenin from the so-called destruction complex that targets β-catenin towards ubiquitin-dependent degradation; free β-catenin then moves to the nucleus and promotes transcription of Wnt targeting genes^57^. GSK3β inhibitors have been widely used to activate Wnt signaling because glycogen synthase kinase (GSK3β) phosphorylates and sets β-catenin for degradation. Here, the selective GSK3 inhibitor SB216763 partially reversed the downregulation of *Syn* in *Abcd* silenced neurons, with no effect in control ones (Fig. 9c.) This finding lends credence to the notion that a deficit in Wnt/β-catenin signaling may contribute to the aberrant development of neurons in X-ALD-like conditions.

In short, the qPCR validations confirm the existence of two sources of damage to X-ALD neurons: cell autonomous mechanisms, shown when ACM^CRTL^ was used, and astrocyte-related ones, manifested in the presence of ACM^X-ALD^. The data also point to different signaling pathways in the autonomous *vs* astrocyte-dependent mechanisms, the former involving members of the Wnt family and the latter mostly Stat3 and Ntrk2 dependent pathways.

We also performed qPCR-based screens in *Abcd1/2* deficient astrocytes to gain insight into molecular transformations rendering ACM^X-ALD^ less permissive for neuronal survival and spinogenesis. We focused on two sets of genes. First, on DEG related to neurodevelopmental pathways detected in the CCALD transcriptomes (Table 1). Even though neuronal development has been more studied than that of astrocytes, some of the dysregulated genes are also critical for astrocytic specification (e.g., *Stat3*^58^ and *Wnt*^57^, reviewed in^59^.) Thus, *a priori* there is no reason to assume that neurons are the only contributors to the differential gene expression detected in whole-brain tissue from CCALD patients. Second, we also validated DEG encoding synaptogenic proteins produced by astrocytes found dysregulated in CCALD transcriptomes (Suppl. data 7.) Synaptogenic proteins include thrombospondins (*Thbs*), glypicans (*Gpc*), ephrins (*Ephb*) and *Sparcl* proteins and, again, members of the Wnt family (reviewed in^60^.)

We found decreased expression of *Wnt2* and *Wnt6*, which were almost completely obliterated, as well as of *Thbs1, Thbs2, Gpc1, Gpc4, Ephrb4* and *Sparcl1*, but not of *Ephrb2, Stat3, Notch3, Nrtk2, Nrtk3*, in X-ALD astrocytes (Fig. 10a). That is, the alterations in *Abcd1/2* deficient astrocytes appear to be relatively selective for pathways involved in synaptogenesis, in accordance with the previous finding of ACM^X-ALD^ mostly affecting spinogenesis among morphometric readouts.

**Figure 10.**
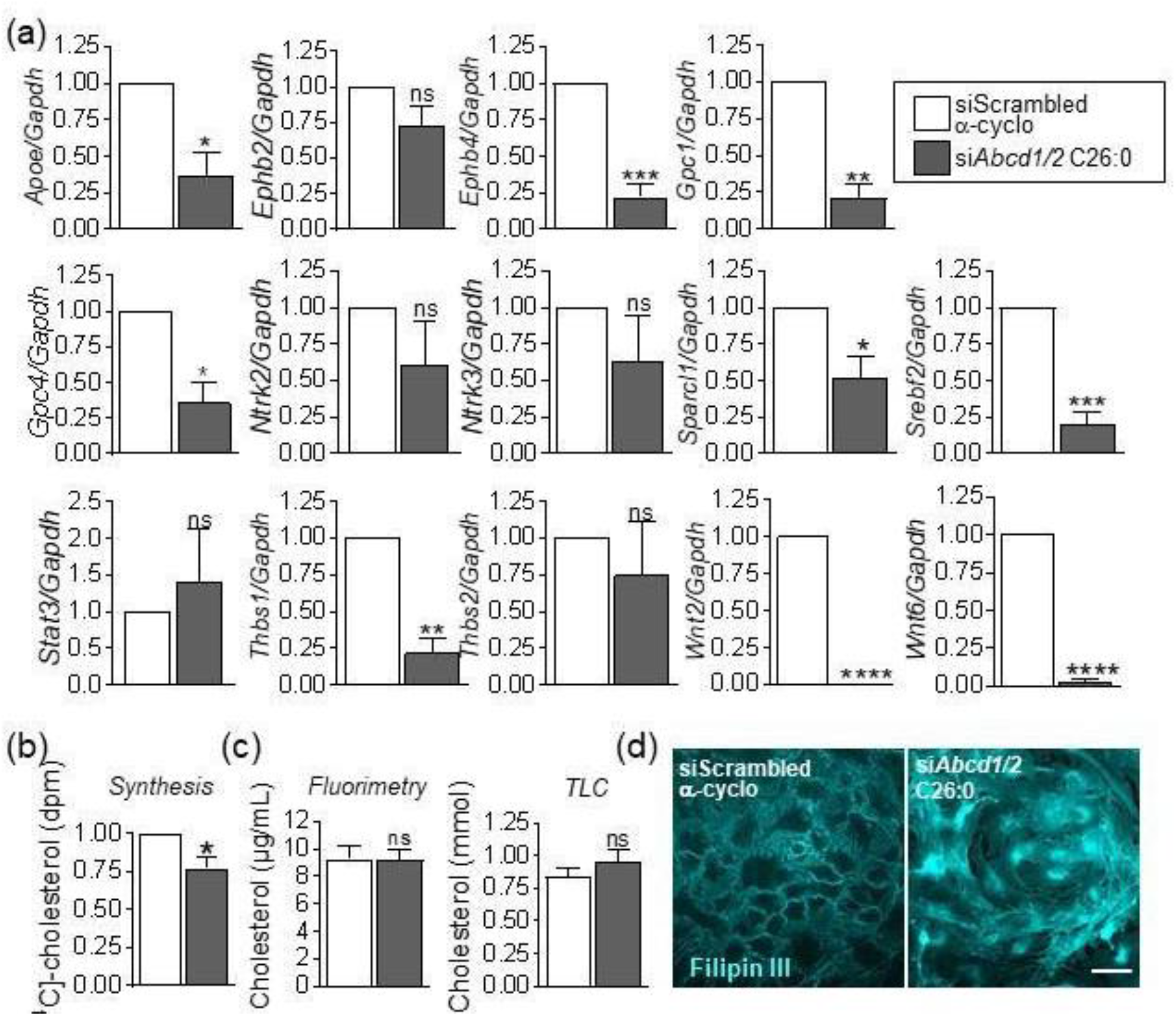
Validation of neurodevelopmental DEG in CCALD in *Abcd1/2* deficient astrocytes. (a) Quantification of mRNA contents of DEG from Table 1 and Suppl. data 7 in control (siScrambled/ α-cyclo) and X-ALD-like astrocytes (*siAbcd1/2* + C26:0) by qPCR. *Gapdh* was used to normalize mRNA data. Data are the means ± SEM; n=3; Unpaired T-test; *p < 0.05; **p < 0.01; ***p < 0.001, ****p<0.0001. *ApoE:* Apolipoprotein E; *Ephb2/4:* Ephrins 4; *Gpc1/4:* Glypicans 1/4; Neurotrophic receptor tyrosine kinase types 2 and 3; *Stat3:* Signal transducer and activator of transcription 3; *Shh:* Sonic hedgehog signaling molecule; *Sparcl1:* Sparc-like protein 1; Srebf2: Sterol Regulatory Element Binding Transcription Factor 2; *Wnt2/ó:* Wnt family members 2 and 6; *Thbs1/2:* Trombospondins1/2. Expression of genes encoding synaptogenic factors is decreased in X-ALD-like astrocytes. (b) Decreased cholesterol biosynthesis as measured by the conversion of the ^14^C-acetate precursor into cholesterol in X-ALD-like conditions. Data are the means ± SEM; n=3; Unpaired T-test. *p < 0.05. (c) Measurement of extracellular cholesterol in astrocyte-conditioned media (ACM) by thin-layer chromatography (TLC) and fluorimetry. Data are the means ± SEM; n=3. No differences were observed in ACM^CRTL^ *vs* ACM^X-ALD^. (d) Altered distribution of cholesterol stained with Filipin III in X-ALD-like astrocytes. Scale bar is 25 μm.

In addition, the expression of *Srebf2* and *ApoE* was decreased in *Abcd1/2* silenced astrocytes (Fig. 10a), which might suggest altered production and delivery of cholesterol, the first astrocytic synaptogenic factor ever to be discovered^61^. Immature neurons produce enough cholesterol to survive but require delivery of cholesterol from astrocytes to form functional synapses^61^. SREBF2 is a transcription factor that promotes synthesis of cholesterol, which is then secreted in ApoE containing lipoproteins that deliver cholesterol to neurons^62^. It is worth noting that cholesterol-related genes are globally dysregulated in CCALD and CAMN according to KEGG and Reactome classifications (Suppl. data 4 and 5), as well as to Gene Set Variation Analysis (GSVA) of LDL-cholesterol transport genes^63^ (Suppl. data 8).

Considering this evidence, we assessed whether the production and secretion of cholesterol was reduced in X-ALD-like astrocytes. The production was reduced, when measured by ^14^C-acetate metabolism (Fig. 10b), but the secretion was not, as shown by the similar cholesterol contents in the ACM from control and silenced cells detected by TLC and fluorimetry (Fig. 10c.) The paradox might be explained by preliminary evidence obtained with the marker of free cholesterol, Filipin III, suggesting alteration of the intracellular distribution cholesterol, which showed frequent localization in clumps in *Abcd1/2* deficient astrocytes, while in control cells it accumulated in plasma membranes in a characteristic beehive pattern (Fig. 10d.) Whether the reduction in cholesterol synthesis is adaptive to prevent aberrant cholesterol accumulation in astrocytes, with no impact on release, remains to be determined, taking into account the complexity of cholesterol homeostasis and fluxes, for the lipid exists in different species (e.g., free vs esterified), intracellular compartments, pools or is lipoprotein bound. For now, the data suggest that global cholesterol dyshomeostasis in X-ALD might implicate astrocytes but do not support a deficit in cholesterol in ACM^X-ALD^. Rather, we attribute the exacerbation of synaptogenesis in *Abcd1/2* silenced neurons incubated in AMC^X-ALD^ to deficits in proteinic synaptogenic factors (Fig. 11.)

**Figure 11.**
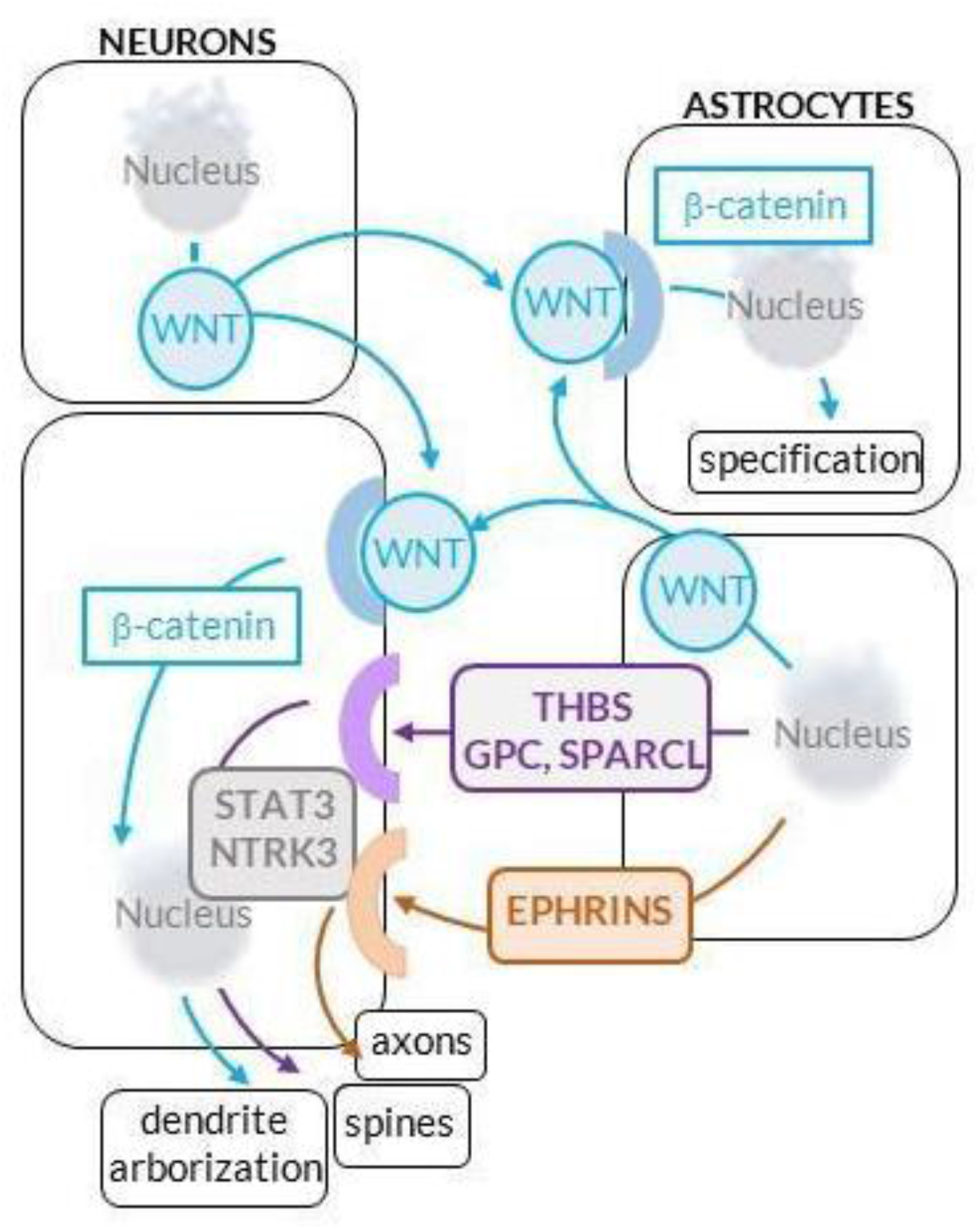
Working model of dysregulated astrocyte-neuron interactions during brain development of carriers of mutated *ABCD*. The model depicts pathways we posit are dysregulated—mostly downregulated— in X-ALD neurodevelopment based on our data from neonatal Abcd*1/2* silenced neurons and astrocytes. In normal brains, WNTs are secreted and act in short ranges as they are hydrophobic, acting either in an autocrine manner, or between functionally and anatomically associated neurons and astrocytes. WNTs trigger neuritogenesis – a generic term defining the formation of spines, axons and dendrite arbors – via beta-catenin mediated transcription; in astrocytes, WNTs trigger specification; synaptogenic factors are then released from astrocytes such as ephrins, trombospondins (THBS), glypicans (GPC) and SPARCL1. The signaling whereby synaptogenic factors from astrocytes promotes neuritogenesis is not known^71^, although the downregulation of *Stat3* and *Nrtk3* in neurons exposed to ACM^X-ALD^ suggests that these pathways are involved in the interactions between astrocytes and neurons during development exists. Plausibly, a certain signaling specialization exists, such that some pathways are more involved in axonogenesis, others in synaptic maturation and others in dendrite growth. In X-ALD, we posit that neurodevelopmental alterations are caused by intrinsic neuronal dysregulation of WNT signaling triggered by aberrant VLCFA accumulation, exacerbated by a deficit of synaptogenic factors, perhaps also caused by impaired Wnt-mediated specification of astrocytes. The nuclei were created with BioRender.com.

## Discussion

We searched for altered neurodevelopmental pathways in brain transcriptomes from childhood and adult cases of cerebral X-ALD to find support for the hypothesis of abnormal neurodevelopment in X-ALD caused by the congenital loss of ABCD function. Next, we performed morphological and molecular validations *in vitro* using a model of early postnatal neuron development in X-ALD based on *Abcd1/2* silencing with RNA interference. Brain development continues postnatally, as shown in rats, the species used in here^64^. In performing the transporter silencing in neurons postnatally we could study the direct consequences of Abcd1/2 malfunction on neuronal growth and development, because any observed change had to be autonomous to the very neurons and not due to damage inflicted on neurons by non-neuronal cells during prenatal stages. For example, the absence of microglia and oligodendrocytes in the cultures allowed us to rule out contributions from microglial secreted factors, free radicals, synaptic pruning and oligodendrocyte degeneration. Finally, our model permitted the analysis of the influence of astrocytes with the widely used ACM approach^65^ that has helped to reveal detrimental actions of astrocytes in neuronal models of acute^49^ and chronic brain diseases^48^. The study of postnatal neuron-astrocyte interactions is moreover appropriate because, in rodents, most of the important waves of synaptogenesis are postnatal and depend on astrocytes^20^. Below we highlight main findings and therapeutic implications.

### First, the greatest disruption of neuronal growth and neuritogenesis in vitro, including axonogenesis, was caused by Abcd1/2 silencing per se, which also rendered neurons dependent on ACM for survival

This finding supports the existence of a cell-autonomous component in neuronopathy in X-ALD^20,21^. Two scenarios are possible: either the accumulation of endogenous VLCFA in *Abcd1/2* silenced neurons and astrocytes is sufficient to induce cellular damage, and/or *Abcd* 1 and 2 have other functions in addition to transporting VLCFA into peroxisomes. For example, a genome-wide shRNA screen for human LDL-cholesterol transport genes has identified *ABCD1* among the relevant genes for cholesterol import into lysosomes in HeLa cells^63^. The relationship between ABCD and cholesterol catabolism may explain why cholesterol-related pathways are dysregulated in subjects harboring *ABCD1* mutations according to the brain transcriptomes.

### Second, dysregulation of specific signaling pathways, namely Wnt signaling, may underlie the disrupted development of Abcd1/2 deficient neurons

This scenario is supported by two pieces of evidence in *Abcd1/2* deficient neurons: the downregulation of *Wnt2* and *Wnt6* but not of genes related to Notch, Shh and neurotrophin signaling, and the rescue of *Syn* expression upon activation of Wnt signaling by a GSK3 pharmacological inhibitor. Wnts are key regulators of several steps involved in dendritic arbor formation such as extension, branching, as well as in stabilization of synaptogenesis through activation of genes controlling actin polymerization^66,67^. Why VLCFA accumulation would affect Wnt signaling can be explained considering some features of Wnt regulation. For example, the fact that fatty acylation of Wnts with C16:0 palmitoleic acid is essential for Wnt function^68^, together with the disruption of lipid homeostasis caused by C26:0^69^, suggest that Wnt may be altered in X-ALD conditions upon defective lipidation. On the other hand, Wnt/beta-catenin signaling is abrogated by oxidative stress, a hallmark of X-ALD^70^, *via* the thioredoxin-like protein nucleoredoxin (NRX)^70^, thus pinpointing another plausible link between Wnt-signaling inhibition and X-ALD pathogenic cascades.

### Third, loss of astrocytic synaptogenic factors may exacerbate abnormal development of Abcd1/2-deficient neurons

Although largely permissive for neuronal growth, ACM^X-ALD^ was not normal, as defective spinogenesis was mildly exacerbated, and the expression of *Stat3* and *Nrtk3* greatly inhibited, in *Abcd1/2* deficient neurons grown in ACM^X-ALD^. It is well documented that ACM contain factors that trigger synaptogenesis and synapse maturation by controlling synaptic vesicle docking, AMPA-receptor clustering, and pre- and post-synapse specialization though trans-synaptic contacts (reviewed in^71^.) The *Abcd1/2*-deficient astrocytes from which ACM^X-ALD^ were collected presented a global downregulation of genes encoding synaptogenic factors, concomitant to a pronounced decrease in *Wnt2* and *Wnt6* expression. Because Wnts elicit astrocyte specification^59^, and because the expression of astrocytic synaptogenic factors is tightly controlled during development in sync with synaptogenesis^71^, the data suggest that *Abcd1/2* deficiency might halt Wnt-elicited astrocyte maturation, thereby canceling synaptogenic cascades. Thus, the altered signaling pathways in the cell-autonomous vs astrocyte-dependent components of abnormal neurodevelopment in mutated ABCD carriers, although distinct, might both involve deficits in Wnt signaling at some stage. In Fig. 11 we integrate these observations in a working model of altered neurodevelopmental astrocyte-neuron interactions in X-ALD to help guide future studies.

### Fourth, limitations of the in vitro models supporting our predictions

One is that neuronal silencing was performed using viral-vectors that *per se* induced neuronal death, which, however small, may render neurons more susceptible to damage caused by subsequent silencing of *Abcd*1/2 transporters. Viral-vector independent approaches to assess neuron-autonomous impaired development are thus in order, for example, are neural stem cells generated from iPSC derived from X-ALD patients abnormal? Another limitation is the use of neurons and ACM from different brain regions for reasons explained earlier. Although this approach arguably unravels basic rules of astrocyte-neuron interactions in X-ALD development, future research should investigate region-specific disturbances. Finally, gene-expression data should be completed with analysis of protein expression in ACMs and functional studies of neuronal activity and plasticity.

### Fifth, ontology and pathway analysis of transcriptomes revealed the same pattern of altered expression of neurodevelopment and neurotransmission pathways in CCLAD and CAMN

The similarity indicates that, despite the profound transformations of the brain during life, the abnormal development leaves a mark in mature neural circuits. However, no gross anatomical alterations have been reported in brains of mutated ABCD1 carriers^30^ and in mouse models with deletions of *Abcd1*^72,73^ and both *Abcd1* and *Abcd2*^40^, although high resolution cytoarchitectural analyses using advanced electron microscopy techniques, or functional connectomics with resting-state functional MRI (rfMRI), have not been used, as MRI remains the gold standard to confirm anatomical abnormalities, namely demyelination, in cerebral forms of the disease. Precedents however exist of alteration of brain circuits in the absence of gross anatomical malformations. This is the case of HD^74^, a neurodegenerative disease with a recently described neurodevelopmental component^26–28^. Of great clinical interest is the capacity of the drug ixoxazole 9 (ISX-9), a small molecule inducer of neurogenesis, neural-cell differentiation and dendrite morphogenesis, to improve cognition and rescue synapse deficits in adult HD mice^27^. These data are important because they causally link abnormal neurodevelopment with pathological manifestations of a neurogenetic disease in adults; how, is not clear. It was argued that *‘subclinical transcriptional alterations could create a different starting point in HD relative to non-HD brain’* in responses to injury^27^, as well as cause mild cognitive impairment in the absence of motor symptoms^37^. Extending these scenarios to X-ALD, cognitive reserve and circuit plasticity may confer resilience against VLCFA accumulation, but perhaps not total protection, as mild psychiatric symptoms and below average performance in some cognitive tests are not rare in adults harboring *ABCD1* mutations, even in ‘pure AMN’ with no MRI abnormalities, all in all pinpointing abnormal brain circuitries in AMN^32–36^. Subclinical or mild neural-circuit dysfunction may accelerate homeostasis loss in injury, hamper repair as well as exacerbate dysfunction in *ABCD1* deficient glia, for glia responses to injury are under the control of neuronal networks^86,88^. If these scenarios are correct, neural-circuit dysfunction due to abnormal development may be considered as another risk factor in X-ALD^16–19^.

### Sixth, therapeutic implications

Current X-ALD therapeutics consist of steroid replacement therapy for the treatment of adrenocortical insufficiency, and hematopoietic stem cell transplantation (HSCT) to arrest CCALD^12^. HSCT, which is thought to replace malfunctional microglia by myeloid cells, is only effective if performed early, when MRI scans show white matter lesions but there are no clinical symptoms^12^. Although newborn screening for X-ALD has been more widely adopted, permitting regular surveillance by MRI and rapid interventions in CCALD, the mortality and morbidity after HSCT remain significant^12^. More recently, gene therapy using patient-derived bone marrow stem cells, based on a seminal study from 2009^76^, was designed as ‘breakthrough therapy’ by the U.S. Food and Drug Administration (FDA), and is being tested in a clinical trial (NCT01896102)^77^. There is not yet treatment for axonal degeneration in AMN, but several therapies are in clinical trials. Two of them, a combination of antioxidants (NCT01495260)^78^, and the PPAR-gamma agonist MIN-102 (NCT03231878), are based on a wealth of evidence in human samples and animal models supporting oxidative stress and mitochondrial deficits as drivers of axonal degeneration in X-ALD (reviewed in^79^). MIN-102 was designated an ‘orphan drug’ by the European Medicines Agency in 2016 and by the FDA in 2017, and a ‘fast drug’ by the FDA in 2020. MIN-102 is also being tested in CCALD before HCTS (2019-000654-59). Finally, there are several trials aimed to establish whether up-regulation of ABCD2 compensates for the lack of ABCD1 (NCT03196765, NCT02559830, NCT03727555). Should future research settle that X-ALD has a neurodevelopmental component, therapeutic strategies under development to correct circuit function and promote neuronal plasticity and tissue repair in neurodevelopmental, neuropsychiatric and neurological disorders might be repurposed for X-ALD; for example, modulators of Wnt signaling^80,81^, trophic factors^82,83^, brain stimulation^84,85^, and epigenetics^88^, to cite a few. Modulation of epigenetic switches has already been considered as a therapeutic avenue in X-ALD^39,89^. Needless to say, the use of multimodal combinations of therapies with distinct cellular and signaling targets will arguably be the most efficacious strategy in X-ALD therapeutics.

## Acknowledgements

This project was partially funded by the European Leukodystrophy Association (ELA2012-033C1) to EG. Current addresses: LP, Cancer Research UK Beatson Institute. CMM: Department of General Pediatrics, Neonatology and Pediatric Cardiology, University Children’s Hospital, H. Heine University, Düsseldorf, Germany.

## Supplementary data

**Suppl. data 1. Gene-expression data used in this study.** There are individual tables for the comparisons between CCALD and AMN with their respective controls. *Excel file attached.*

**SUPPL. DATA 2.**
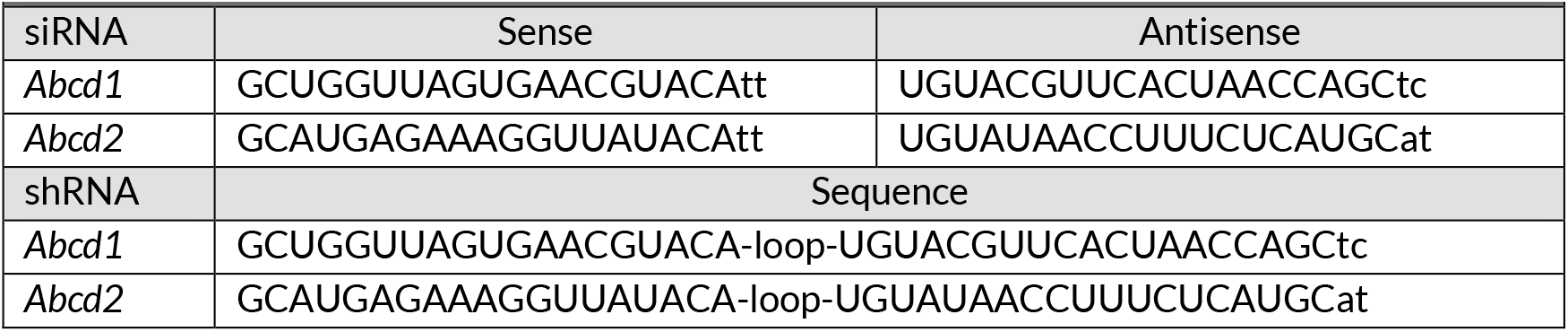
Sequences of siRNAs and shRNAs for Abcd transporters.

**SUPPL. DATA 3.**
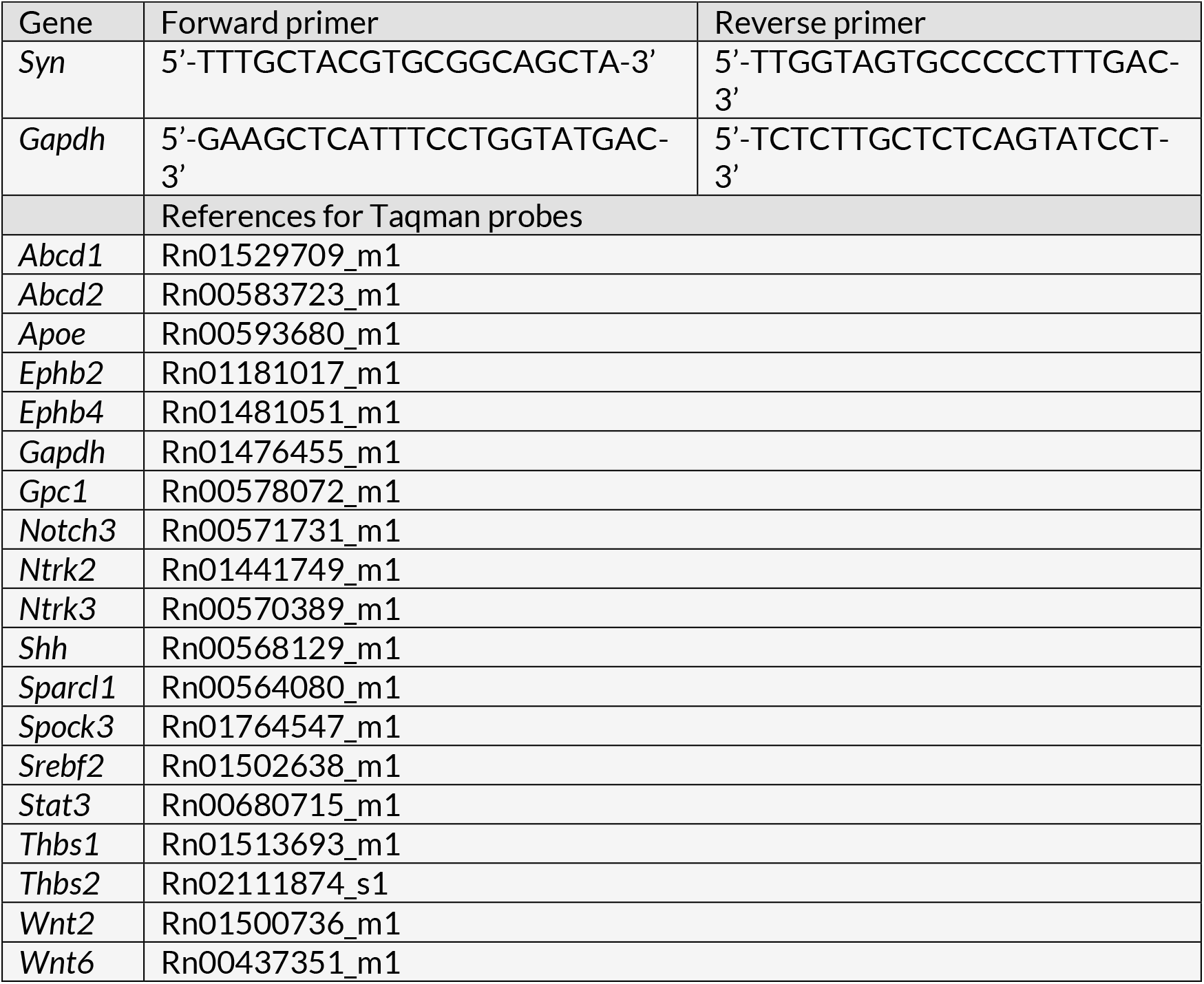
Primer sequences.

**SUPPL. DATA 4.** Pathway dysregulation CNS development/neurotransmission/apoptosis CCALD vs Controls.

|  |  |  |  |  |  |
| --- | --- | --- | --- | --- | --- |
| Platform |  | Rank | Overlap | p-values | Adj p-values |
| REACTOME<br>Human<br>2016 | Developmental biology<br>Homo sapiens R-HSA-1266738 | 3 | 221/786 | 1.67E-12 | 8.25E-10 |
|  | Axon guidance Homo Sapiens R-HSA-422475 | 16 | 144/515 | 1.97E-08 | 1.82E-06 |
|  | Neuronal System Homo Sapiens R-HSA-112316 | 36 | 90/301 | 3.95E-07 | 1.62E-05 |
|  | Programmed Cell Death<br>Homo sapiens R-HSA-5357801 | 45 | 55/166 | 2.42E-06 | 7.94E-05 |
|  | Transmission across chemical<br>Synapses<br>Homo sapiens R-HSA-112315 | 59 | 64/211 | 1.07E-05 | 2.61E-04 |
|  | Regulation of cholesterol<br>biosynthesis by SREBP (SREBF)<br>Homo sapiens R-HSA-1655829 | 61 | 24/55 | 1.08E-05 | 2.61E-04 |
| GO_BP<br>2021 | Anterograde trans-synaptic signaling<br>(GO:0098916) | 7 | 80/244 | 2.38E-08 | 1.90E-05 |
|  | Regulation of apoptotic process<br>(GO:0042981) | 8 | 187/742 | 6.56E-07 | 3.07E-04 |
|  | Axon guidance (GO:0007411) | 23 | 63/203 | 5.55E-06 | 0.00135322 |
|  | Apoptotic process (GO:0006915) | 25 | 69/231 | 8.85E-06 | 0.00198579 |
|  | Axonogenesis (GO:0007409) | 66 | 67/240 | 1.20E-04 | 0.01021901 |
|  | Nervous system development<br>(GO:0007399) | 70 | 112/447 | 1.41E-04 | 0.01131636 |
|  | Central nervous system development<br>(GO:0007417) | 86 | 72/268 | 2.47E-04 | 0.01569497 |
|  | Positive regulation of cell migration<br>(GO:0030335) | 93 | 72/269 | 2.79E-04 | 0.01667884 |
| GO_CC<br>2021 | Dendrite (GO:0030425) | 3 | 84/270 | 1.48E-07 | 2.32E-05 |
|  | Neuron projection (GO:0043005) | 4 | 147/556 | 5.86E-07 | 6.87E-05 |
|  | Axon (GO:0030424) | 15 | 58/204 | 1.97E-04 | 0.005841492 |
| KEGG_<br>Human<br>2021 | Neuroactive ligand-receptor<br>interaction | 2 | 121/341 | 1.29E-14 | 2.06E-12 |
|  | Apoptosis | 99 | 41/142 | 0.001138 | 0.00366952 |
|  | Axon guidance | 101 | 50/182 | 0.001202 | 0.00379931 |
|  | Cholesterol metabolism | 150 | 16/50 | 0.012855 | 0.02733999 |
| Pathway (Reactome, KEGG) and ontology (GO-CC and GO-BP) analyses of DEG of CCALD showing categories relevant to CNS development with adjusted p-values <0.05. Analysis performed with Enrichr. |  |  |  |  |  |

**SUPPL. DATA 5.** Pathway dysregulation CNS development/neurotransmission/apoptosis AMN vs Controls.

| Platform |  | Rank | Overlap | p-values | Adj p-values |
| --- | --- | --- | --- | --- | --- |
| REACTOME<br>Human<br>2016 | Developmental biology<br>Homo sapiens R-HSA-1266738 | 5 | 280/786 | 9.81E-13 | 2.95E-10 |
|  | Axon guidance<br>Homo sapiens R-HSA-422475 | 10 | 187/515 | 1.12E-09 | 1.69E-07 |
|  | Transmission across biochemical<br>Synapses Homo sapiens<br>R-HSA-112315 | 66 | 75/211 | 2.18E-04 | 0.0049663 |
|  | Neuronal System *Homo sapiens*<br>R-HSA-112316 | 188 | 96/301 | 0.0022298 | 0.0178385 |
|  | Regulation of cholesterol<br>biosynthesis by SREBP (SREBF)<br>Homo sapiens<br>R-HSA-1655829 | 226 | 23/55 | 0.0035858 | 0.0237582 |
| GO_BP<br>2021 | Regulation of cell migration<br>(GO:0030334) | 16 | 154/408 | 1.48E-09 | 5.35E-07 |
|  | Regulation of apoptotic process<br>(GO:0042981) | 18 | 252/742 | 2.60E-09 | 8.45E-07 |
|  | Apoptotic process (GO:0006915) | 26 | 94/231 | 3.89E-08 | 8.74E-06 |
|  | Positive regulation of cell migration<br>(GO:0030335) | 40 | 58/144 | 2.09E-05 | 0.0019052 |
|  | Nervous system development<br>(GO:0007399) | 41 | 154/447 | 1.26E-06 | 1.80E-04 |
|  | Central nervous system development<br>(GO:0007417) | 47 | 99/268 | 3.60E-06 | 4.48E-04 |
|  | Axon guidance (GO:0007411) | 67 | 76/203 | 2.72E-05 | 0.0023734 |
|  | Axonogenesis (GO:0007409) | 102 | 85/240 | 9.70E-05 | 0.0055556 |
|  | Anterograde trans-synaptic signaling<br>(GO:0098916) | 220 | 82/244 | 8.38E-04 | 0.0222353 |
| GO_CC<br>2021 | Neuron projection (GO:0043005) | 17 | 185/556 | 1.58E-06 | 4.57E-05 |
|  | Dendrite (GO:0030425) | 42 | 91/270 | 4.06E-04 | 0.0047548 |
|  | Axon (GO:0030424) | 59 | 68/204 | 0.0027689 | 0.0227054 |
| KEGG<br>Human<br>2021 | Neuroactive ligand-receptor<br>interaction | 9 | 128/341 | 5.09E-08 | 1.81E-06 |
|  | Apoptosis | 21 | 60/142 | 2.50E-06 | 3.81E-05 |
|  | Axon guidance | 68 | 66/182 | 2.59E-04 | 0.0012197 |
| Pathway (Reactome, KEGG) and ontology (GO-CC and GO-BP) analyses of DEG of and AMN vs control showing categories relevant to CNS development with adjusted p-values <0.05. Analysis performed with Enrichr. |  |  |  |  |  |

**SUPPL. DATA 6.** Statistics of TUNEL assay in neurons corresponding to Fig. 3B.

|  |  |  |  |  |  |  |  |  |  |  |  |  |  |
| --- | --- | --- | --- | --- | --- | --- | --- | --- | --- | --- | --- | --- | --- |
|  |  | shScrambled Veh |  |  | shScrambled C26:0 |  |  | shAbcd1/2 Veh |  |  | shAbcd1/2 C26:0 |  |  |
|  |  | Minus ACM | CTRL ACM | XALD ACM | Minus ACM | CTRL ACM | XALD ACM | Minus ACM | CTRL ACM | XALD ACM | Minus ACM | CTRL ACM | XALD ACM |
| shScrambled Veh | Minus ACM |  | ns | ns | * |  |  | **** |  |  | **** |  |  |
|  | CTRL ACM | ns |  | ns |  | ns |  |  | * |  |  | ns |  |
|  | X-ALD ACM | ns | ns |  |  |  | ns |  |  | *** |  |  | *** |
| shScrambled C26:0 | Minus ACM | * |  |  |  | *** | **** | **** |  |  | **** |  |  |
|  | CTRL ACM |  | ns |  | *** |  | ns |  | ** |  |  | * |  |
|  | X-ALD ACM |  |  | ns | **** | ns |  |  |  | **** |  |  | **** |
| shAbcd1/2 Veh | Minus ACM | **** |  |  | **** |  |  |  | **** | *** | ns |  |  |
|  | CTRL ACM |  | * |  |  | ** |  | **** |  | ns |  | ns |  |
|  | X-ALD ACM |  |  | *** |  |  | **** | *** | ns |  |  |  | ns |
| shAbcd1/2 C26:0 | Minus ACM | **** |  |  | **** |  |  | ns |  |  |  | **** | *** |
|  | CTRL ACM |  | ns |  |  | * |  |  | ns |  | **** |  | ns |
|  | X-ALD ACM |  |  | *** |  |  | **** |  |  | ns | *** | ns |  |

(\*) p<0.05; (\*\*) p<0.01, (\*\*\*) p<0.001 and (\*\*\*\*) p<0.0001; Two-way ANOVA and post-hoc Tukey's multiple comparisons test; ns: non-significant; squares in grey: non-relevant comparisons.

**SUPPL. DATA 7.**
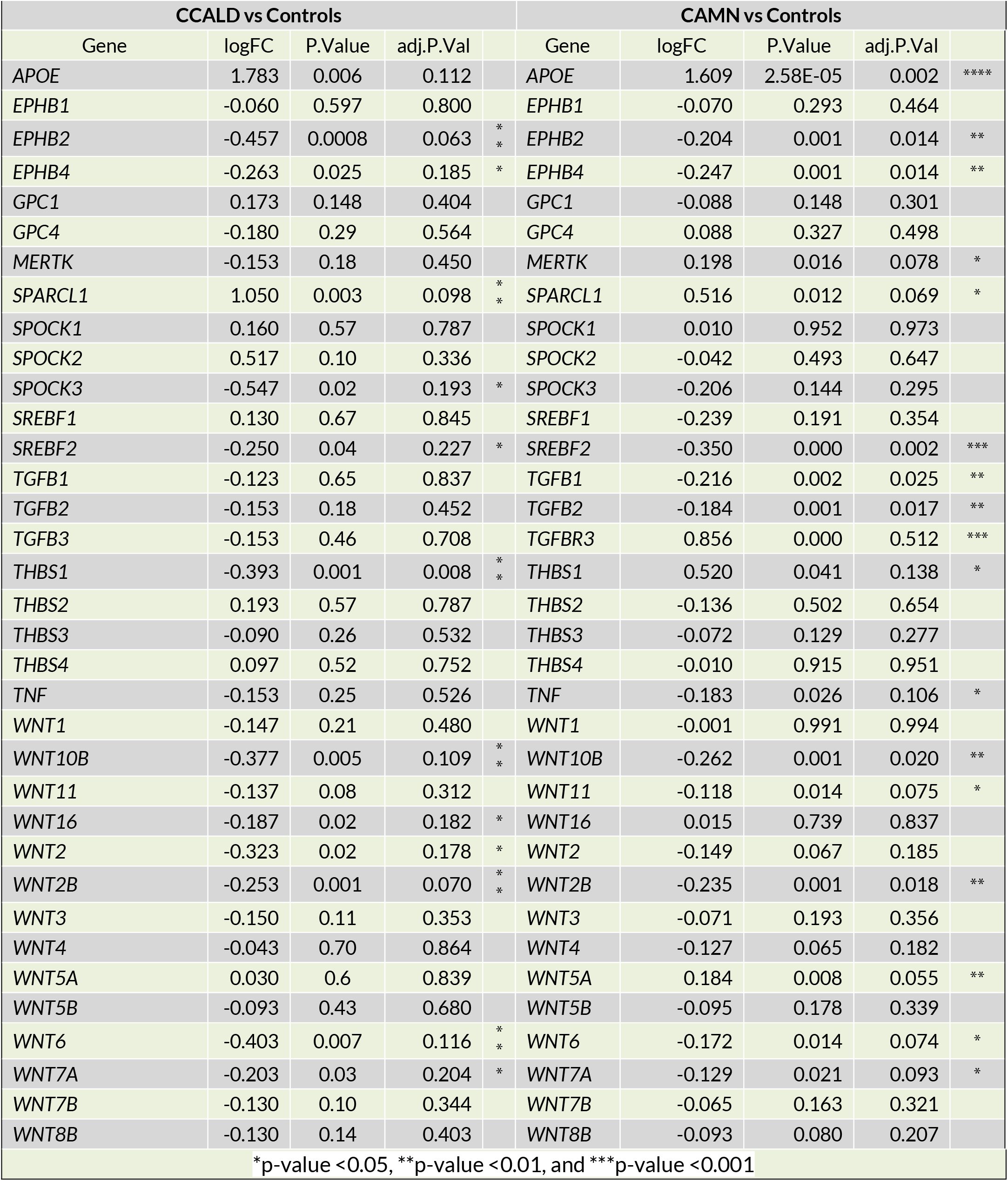
Expression of genes encoding for synaptogenic factors in CCALD and CAMN.

**SUPPL. DATA 8.**
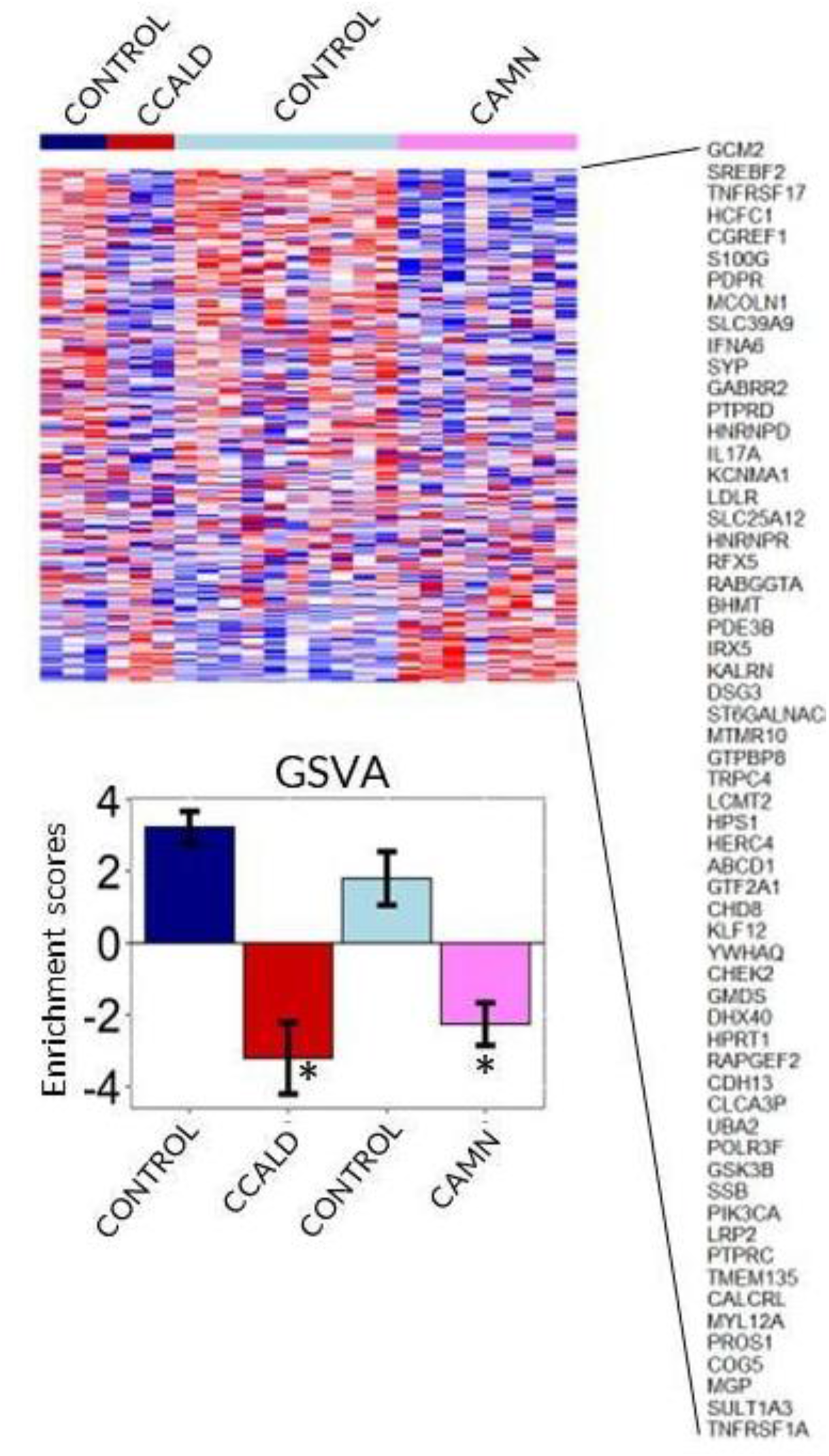
Expression of genes identified in a genome-wide shRNA screen for LDL-cholesterol transport genes^63^ in X-ALD human samples. All samples from all databases (young controls, CCALD, adult controls and CAMN) were analyzed together using Gene Set Variation Analysis (GSVA); Only some genes representative of the 341 identified in^63^ are shown in the right column for clarity. Cholesterol-transport genes are downregulated in CCALD and CAMN as compared to age-matched controls. (*) qFDR<0.05; X-ALD vs Controls.

